# An ancient polymorphism in myosin I a/b determines the left-right asymmetry of Japanese snails

**DOI:** 10.64898/2026.06.09.731057

**Authors:** Alec Lewis, Yasuto Ishii, Shun Ito, Kazuki Kimura, Margrethe Johansen, Alistair Hume, Christopher Moore, Tom Hartman, Daniel J. Jackson, Satoshi Chiba, Takahiro Hirano, Angus Davison

## Abstract

Left-right (LR) asymmetry is a fundamental feature of animal development, yet progress in understanding its establishment has been limited by the fact that almost all animals exhibit invariant chirality. Snails are the exception because some wild-living species show abundant chiral variation, yet the existing knowledge is from snails in which the chirality mutation is rare and causes pathology. Here, we show that a chiral polymorphism in Japanese *Euhadra* snails is due to functional variation in an unconventional myosin I a/b, an isoform not previously implicated in LR specification in any animal. Both the sinistral and dextral gene variants are ancient, impart minimal transcriptional differences in the single-cell embryo and are not pathological. Phylogenetic and structural modelling suggests that dominant-acting amino acid substitutions in the myosin actin-binding domain and/or motor-level junction were enabled by relaxed selection. These results broaden the known molecular repertoire underlying LR asymmetry, suggest key mutations and positions that should be explored in other model animals, and highlight snails as a powerful model for understanding the origins of animal LR asymmetry.

## Introduction

Defining the left and right sides of the body is a critical part of animal development^1-3^, yet a limiting factor for studying asymmetry is that the primary or embryonic left-right (LR) axis is invariant (e.g. heart to the left in humans). In consequence, the biology and genetics of an evolutionary change of the primary axis of LR asymmetry has never been observed in a wild-living population. To further understanding, one experimental method has been to use mutants or manipulate embryos of a select group of model animals to create individuals that are partly or wholly orientated in the opposite direction.^3-5^ A more recent approach has been to understand how chirality can arise from interaction of chiral molecular structures, cortical flow or cells^2,6-9^, and then make links across scales.^10^ These approaches are valuable, but limited, in the sense that they primarily illustrate why chirality does not vary, and what happens when it goes wrong, including in humans.^11^ They do not shed light on why LR asymmetry does sometimes vary and how body-plans change over evolutionary time.

Ironically, scientists have largely ignored the only animal group – snails – in which ordinary development commonly produces individuals that are wholly mirror-imaged, reflected outwardly in right- (dextral) or left-coiling (sinistral) shells.^12^ Famously, chirality in snails is an inherited condition, determined by a single Mendelian locus of maternal effect.^12-14^. The chirality gene product directs a clockwise or anti-clockwise spiral cell cleavage^12,15^ early in development, ultimately producing snails with chirally asymmetric bodies and shells.

Unfortunately, limitations in technologies and access to specimens have meant that prior studies^16-18^ have been limited to using mutant snails in which the mirror-imaging is pathological. Thus, while it was an important finding to show that a frameshift mutation in a duplicated formin gene is responsible for producing the ultra-rare sinistral chiral type in the pond snail *Lymnaea stagnalis*^16,17^, only about 50% of embryos hatch.^19,20^ In comparison, the abundant chiral variation that is present in wild-living land snails has been sifted by natural selection, but the genes that determine this natural variation remain completely unknown. Here, we show that LR asymmetry in Japanese *Euhadra* snails (**Figure 1A**) is associated with functional variation in an unconventional myosin I a/b, an isoform not previously implicated in LR specification in any animal. The findings illustrate the biological and genetic basis of natural variation in snail chirality, and lay the groundwork for a deeper mechanistic understanding of why other animals do not vary in their LR asymmetry.

**Figure 1.**
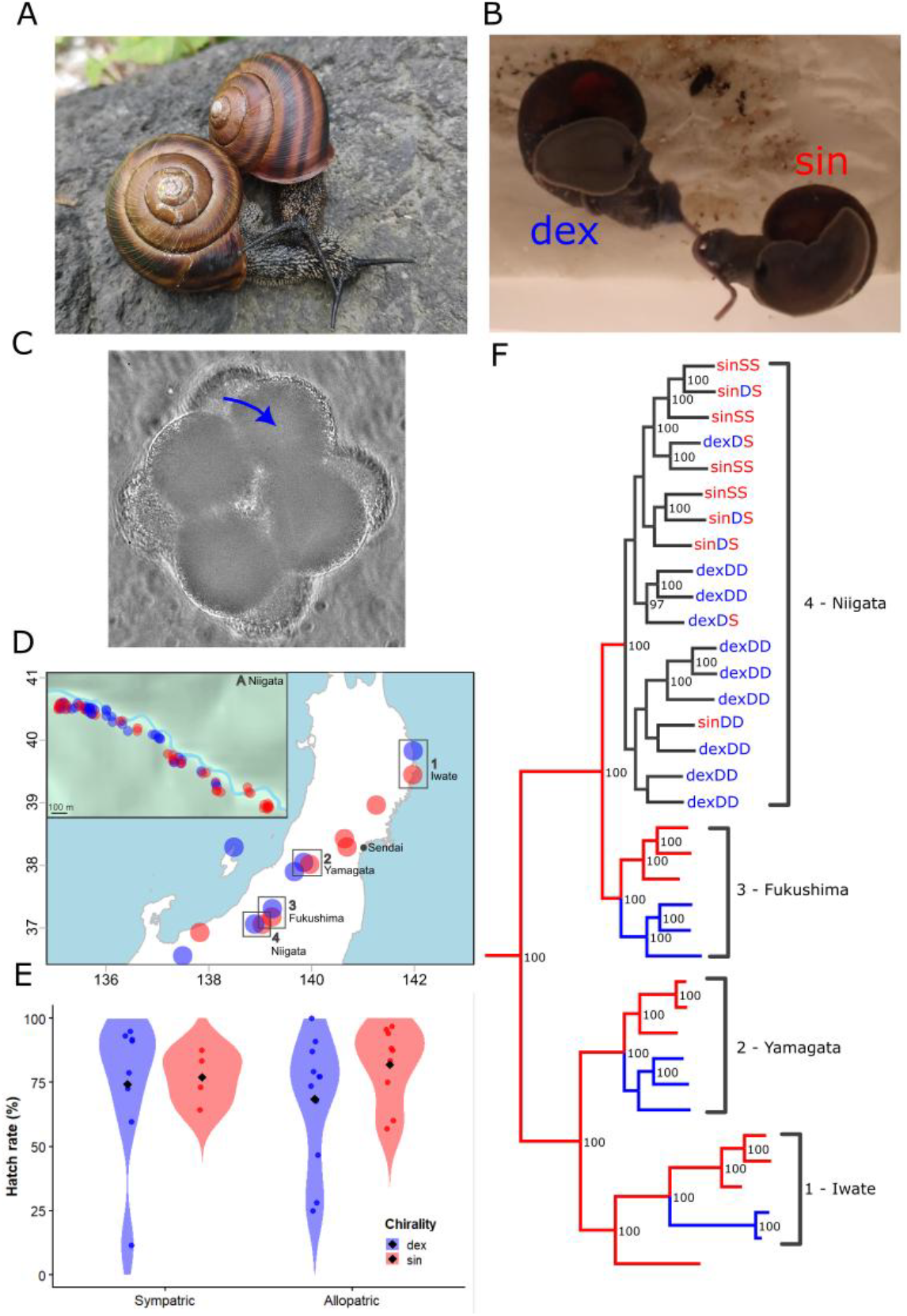
Gene flow between sinistral and dextral snails. **(A)** Sinistral and dextral *E. quaesita* from sympatric site, showing similar shell morphology. **(B)** Unilateral mating between dextral and sinistral *E. quaesita* in laboratory (inverted on glass). Dextral snail has inserted penis, sinistral has not. **(C)** Third cleavage of *E. quaesita* embryo showing dextral twist. **(D)** Main sample locations in northern Japan, showing three sites where dextral (blue) and sinistral (red) *E. quaesita* are allopatric, but physically close to each other (#1 to #3), or sympatric (#4). Inset shows detail of sympatric zone at site #4. **(E)** Egg hatch rates were similar between sinistral and dextral individuals, regardless of whether populations were sympatric or allopatric (see Supplementary Information for statistical tests). **(F)** Whole-genome maximum-likelihood phylogeny focussed on the monophyletic *E. quaesita* group and the main sample sites. Except in Niigata, snail chiral genotype is the same as chiral phenotype, so lineages are coloured either red or blue for sinistral or dextral. At site #4 in Niigata there is present-day gene-flow, so both chirality phenotype (sin/dex) and genotype (*S*/*D*) is shown. Note that the full chirality genotype of these site #4 snails was inferred from the GWAS.

## Results and Discussion

### Gene-flow between dextral and sinistral snails

A main difficulty in identifying a LR asymmetry-determining gene in snails is that dextral and sinistral snails tend to have largely separate distributions because the two types have difficulty mating, and are arguably reproductively isolated by ‘single-gene’ speciation.^21-23^ If there is no gene-flow then a genome-wide association study is not possible. To get around this issue we identified three putative contact zones between dextral and sinistral Japanese *Euhadra quaesita* snails in northern Japan (#1 to #3), and a fourth site in Niigata (#4) where both chiral types are fully sympatric and have shells that look similar (**Figures 1A, D; see Tables S1, S2**).

In the lab, snails from the sympatric site (#4) always produced offspring of a fixed chirality, some of which were of the opposite chirality to the mother (**Table S3**). As expected, this chirality was evident in the chiral twist that takes place during the third cell cleavage in early development (**Figure 1C; Video S1**). Together, these observations indicate that chirality in *E. quaesita* is maternally inherited, as in other snails^12^, and that there must be present-day gene-flow between sinistrals and dextrals at the site in Niigata (#4). Gene-flow is enabled by the maternal inheritance of chirality which means that there is a generational delay in the expression of chirality^22,24^, and more straightforwardly because mating is at least sometimes possible between sinistrals and dextrals (**Figure 1B; Video S2**).

No gross differences were discovered in the hatch rate of eggs from snails that are genetically dextral or sinistral (**Figure 1E; Table 3, Supplementary Results**). The chirality locus in *Euhadra* therefore determines LR asymmetry but otherwise is not pathological, while not ruling out more subtle effects.

A whole genome phylogeny showed that there is a monophyletic group which contains the four wholly sinistral species, as well as chirally variable *E. quaesita* (**Figures 1F, S1**), a result that is consistent with a single evolution of sinistrality, with subsequent reversions to dextrality within *E. quaesita*. Snails at the sympatric site (#4), form a monophyletic group but are otherwise mixed for chirality, which is indicative of present-day gene flow between the two chiral types (**Figure 1F**). Snails from the other three contact zones are on three separate branches, and cluster by chirality. Structure-based analyses clustered *E. quaesita* snails by geography, rather than by chirality (**Figure S2;** see **Figure S3** for similar findings using principal components analysis). At each paired site, differentiation between sinistral and dextral is low in most regions of the genome (**Figure S4**). These patterns are most simply explained by present-day or recent gene flow between sinistrals and dextrals at each of the four sites, with local populations (at sites #1 to #3) tending to fix for one or the other chiral type, due to frequency-dependent selection against the rare type.

### Unconventional myosin I determines LR asymmetry

Genome-wide association methods showed two narrow peaks of association (**Figures 2A, S5**), one on chromosome 27 for which there are multiple statistically significant SNPs (positions 73.6-74.1 Mb), and another on chromosome 4 for which most SNPs (positions ∼17.7-17.9 Mb) were just below the strict statistical threshold (**Figure 2B;** see **Table S4**). Sliding window analyses of Fst showed a consistent signal at the same position on chromosome 27, but not always on chromosome 4 (**Figure S4**), and with low values of absolute divergence (Dxy). These findings are consistent with chiral variation in *Euhadra* being caused by variation at a locus on chromosome 27 between positions 73.6-74.1 Mb, while not ruling out the possible influence of another locus. Chiral evolution may be due to an external selection pressure, such as reproductive character displacement^12^ or chiral predation^25^, but there is no evidence that the two chiral types are in the process of speciating; ‘single-gene’ speciation is discounted.^22,23^

**Figure 2.**
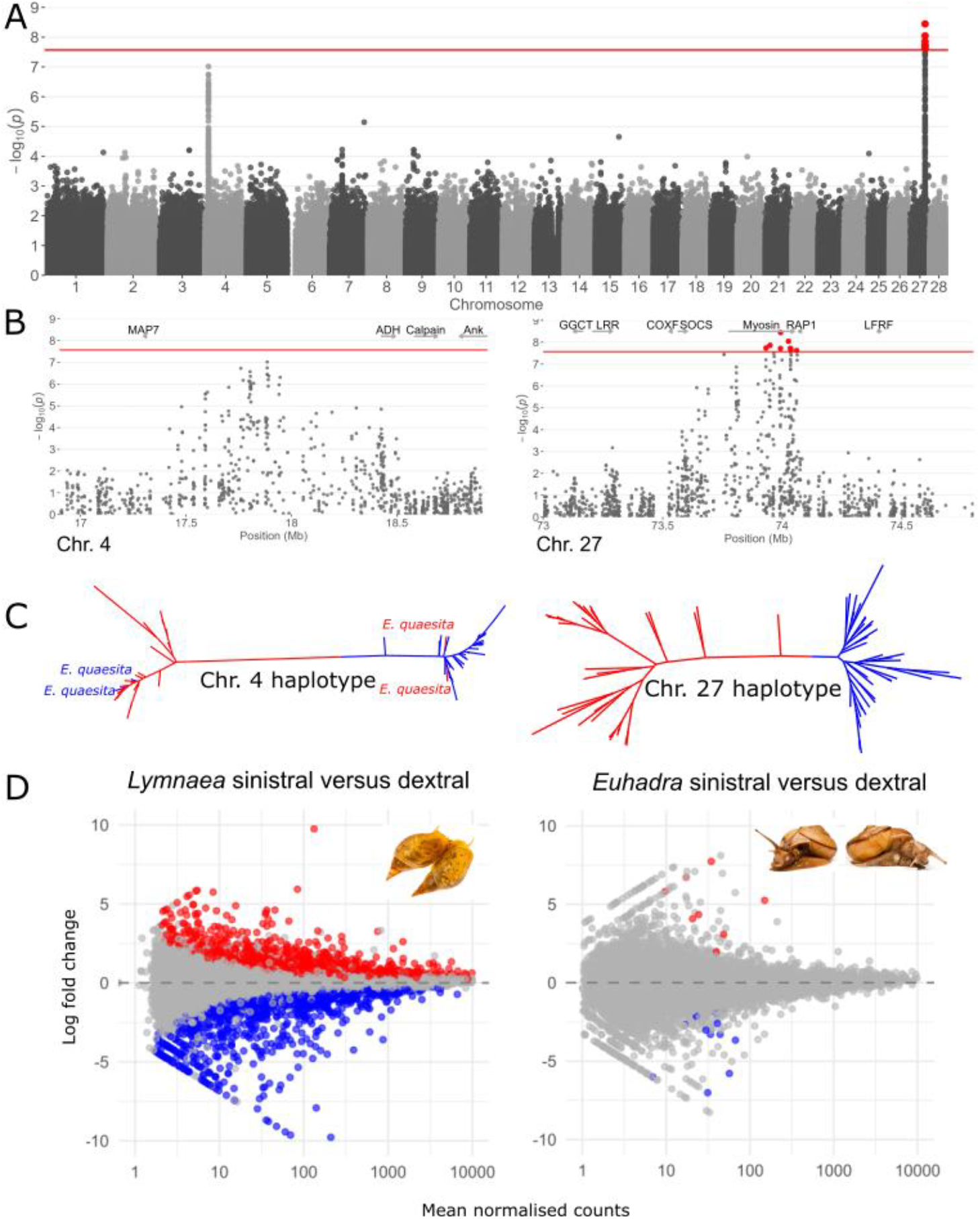
Genomic analyses of sinistral and dextral *Euhadra*. **(A)** Genome-wide association analysis shows two peaks of association, one that breaches significance threshold on chromosome 27 and a lesser peak on chromosome 4. **(B)** The peak on chromosome 27 is directly over a myosin gene, and also RAP1. No genes were discovered that are associated with the peak on chromosome 4. **(C)** Phylogenies produced using individual haplotypes of *E. quaesita*. The phylogeny for the chromosome 27 region perfectly separates dextral (blue) from sinistral (red), the only region in the genome to do so. In comparison, two individuals are anomalous (both homozygotes, E8 and E187; Figure S7) for the chromosome 4 region. **(D)** No differences in myosin expression were discovered between dextral and sinistral *E. quaesita*, with only minimal transcriptional differences in other genes (22 genes, 0.08%). In comparison, 3005 genes (17%) were differentially expressed between the dextral and sinistral pond snail embryos, including the formin gene, *Lsdia2*. Red = upregulated in sinistral, blue = downregulated in sinistral.

Further analyses showed that linkage disequilibrium is generally low on all chromosomes (**Figure S6**), declining to a background rate (*r*^*2*^ ∼ 0.06) once the pairwise distance is around 50-75 kb. Topology-weighting methods on a subset of individuals confirmed that across the whole genome there are only two regions in which the phylogenetic signal tends to group populations by chirality, a region of 462 kb on chromosome 27, and another of 168 kb on chromosome 4 (**Figures S7, S8**). Both of these regions are exactly concordant with the peaks observed in both the GWAS and sliding window Fst analyses.

A phylogeny using individual haplotypes of the region for chromosome 27 showed sinistral *E. quaesita* as monophyletic, consistent with this region containing the chirality-determining gene (**Figure 2C**). In comparison, two individuals were anomalous in a chromosome 4 region phylogeny (**Figures 2C, S8**). These general findings are supported by much wider sampling using an independent dataset with a much larger set of *E. quaesita* and applying ddRAD methods (**Figure S9**). Hence, the chirality locus must be on chromosome 27, between positions ∼73.6-74.1 Mb. An all species phylogeny of the same region (**Figure S10)** shows that both the dextral and sinistral lineages derived from the chromosome 27 region are ancient, and not closely related to any other lineages within the genus. The deep evolutionary history of the polymorphism is also consistent with a lack of pathology (**Figures 3C, S10**).

**Figure 3.**
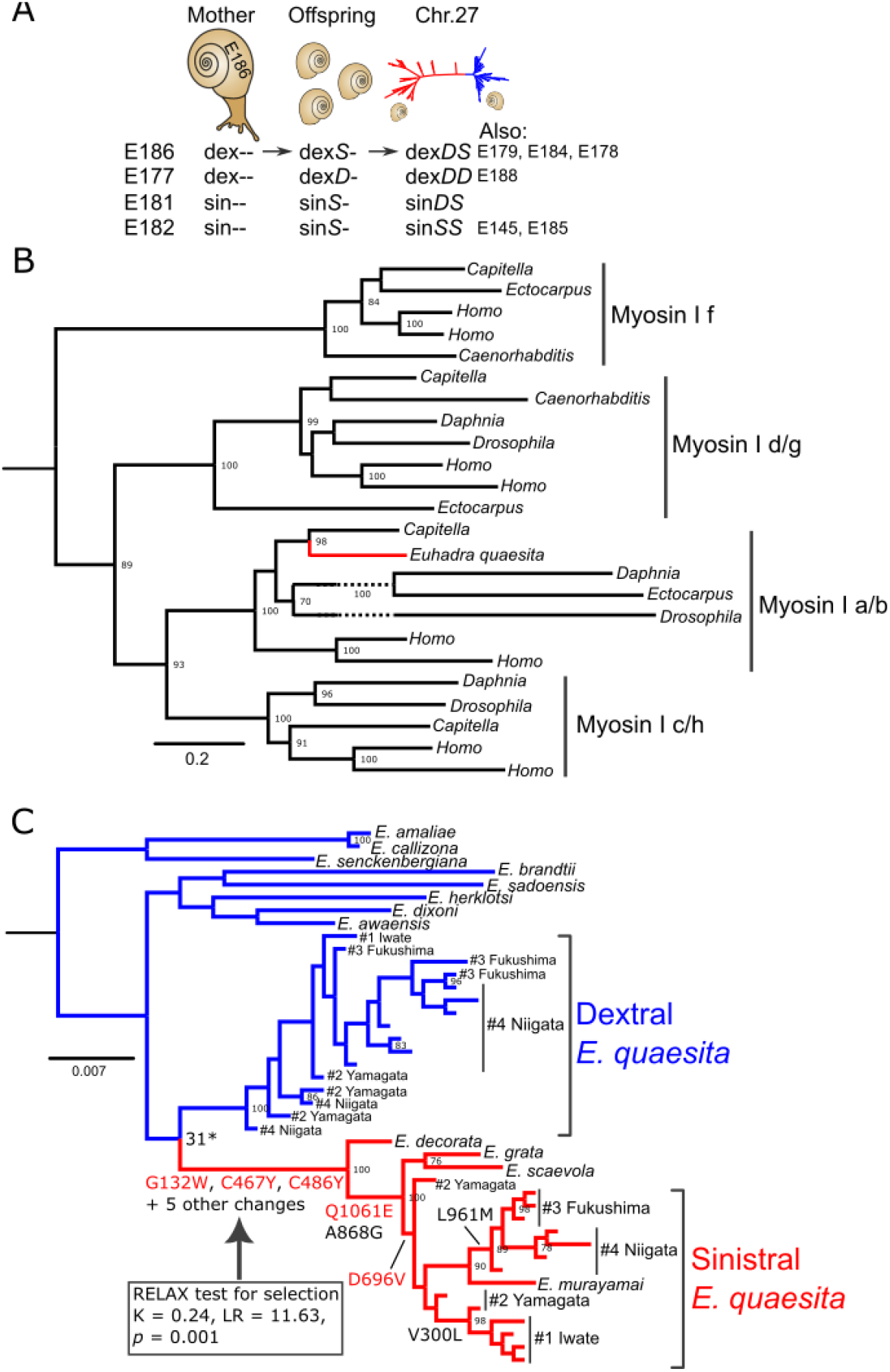
Evolution and inheritance of the myosin gene that determines chirality in *Euhadra*. **(A)** Scheme illustrating inference of dominance, example shows snail E186. A dextral snail (“dex”), E186, produced sinistral offspring so must be genetically sinistral (“dex*S*”). Snail E186 has two very divergent haplotypes at the chr. 27 locus, one that defines dextral and one that defines sinistral. E186 is therefore a dextral snail of genotype *DS*, where *S* is dominant. **(B)** An amino acid-based phylogeny of the motor domain shows that the snail gene is a myosin I a/b isoform. **(C)** Phylogeny of the myosin gene in *Euhadraą* also showing amino acid changes (non-conservative in bold). Sinistral *Euhadra* are monophyletic. There are multiple amino acid changes in the branches that define sinistrality but none at all in the other branches. Statistical tests show evidence for relaxed selection on the sinistral branch. Note that the branch that separates sinistral from dextral (marked with an asterisk) is strongly supported when using the 462 kb chromosome 27 region.

A custom genome annotation and transcriptomic evidence were used to identify genes within the candidate region on chromosome 27, while also inspecting chromosome 4. The SNPs on chromosome 27 are entirely associated with a region that contains an unconventional myosin I gene for which the exons span exactly the same candidate region, 73.6-74.1 Mb. The only other possible candidate is a putative low density lipoprotein receptor adaptor (RAP1) gene (**Figure 2B**), but this is not present as mRNA in single-cell embryos (**Table S5**), and so is not likely involved. No embryo-expressed genes were discovered near the peak on chromosome 4. Thus, all evidence points to the myosin gene as the chirality-determining locus.

Individual haplotypes for the chromosome 27 region were compared with snails of known chirality genotype. These analyses showed that the sinistral allele is dominant-acting, which likely gives some clue as to its function (**Figure 3A**; **Tables S2, S3**).

To further understand chiral variation, RNA-seq methods were applied to individual single-cell embryos, comparing gene expression between sinistral and dextral *E. quaesita*. No differences in myosin expression were discovered between dextrals and sinistrals, with only minimal transcriptional differences in other genes (22 genes, 0.08%), including membrane-associated and zinc-finger genes (**Figures 2D, S11**; **Tables S6, S7**). Chiral variation is thus not likely caused by differential expression of myosin mRNA in the single-cell embryo.

As a point of comparison, the same RNA-seq experiment was carried out using sinistral and dextral embryos of the pond snail for which the causal mutation is a frameshift in the *Lsdia2* formin gene.^16^ ln contrast to *E. quaesita*, 3005 genes (17% of total) were differentially expressed between the two *L. stagnalis* embryo types (**Figure 2D; Tables S8, S9**), including the formin gene, *Lsdia2*, as expected. This therefore further emphasises the issue that investigating a pathological mutant is a poor proxy compared with natural variation.

### LR asymmetry is due to amino acid changes in myosin 1a/b

A phylogeny shows that the myosin gene is an unconventional myosin I a/b (**Figure 3B**), an isoform that is involved in general actin and membrane biology but has not previously been implicated in LR asymmetry, despite intense study in *Drosophila*.^26,27^ A combination of structural modelling and phylogenies indicate that the *E. quaesita* myosin has the canonical motor domain, three IQ motifs immediately terminal to the motor domain and a TH1 tail (**Figure 4A**), including a full motor domain, a multi-IQ lever arm immediately C-terminal to the converter, and a TH1 PH-like tail for membrane association

**Figure 4.**
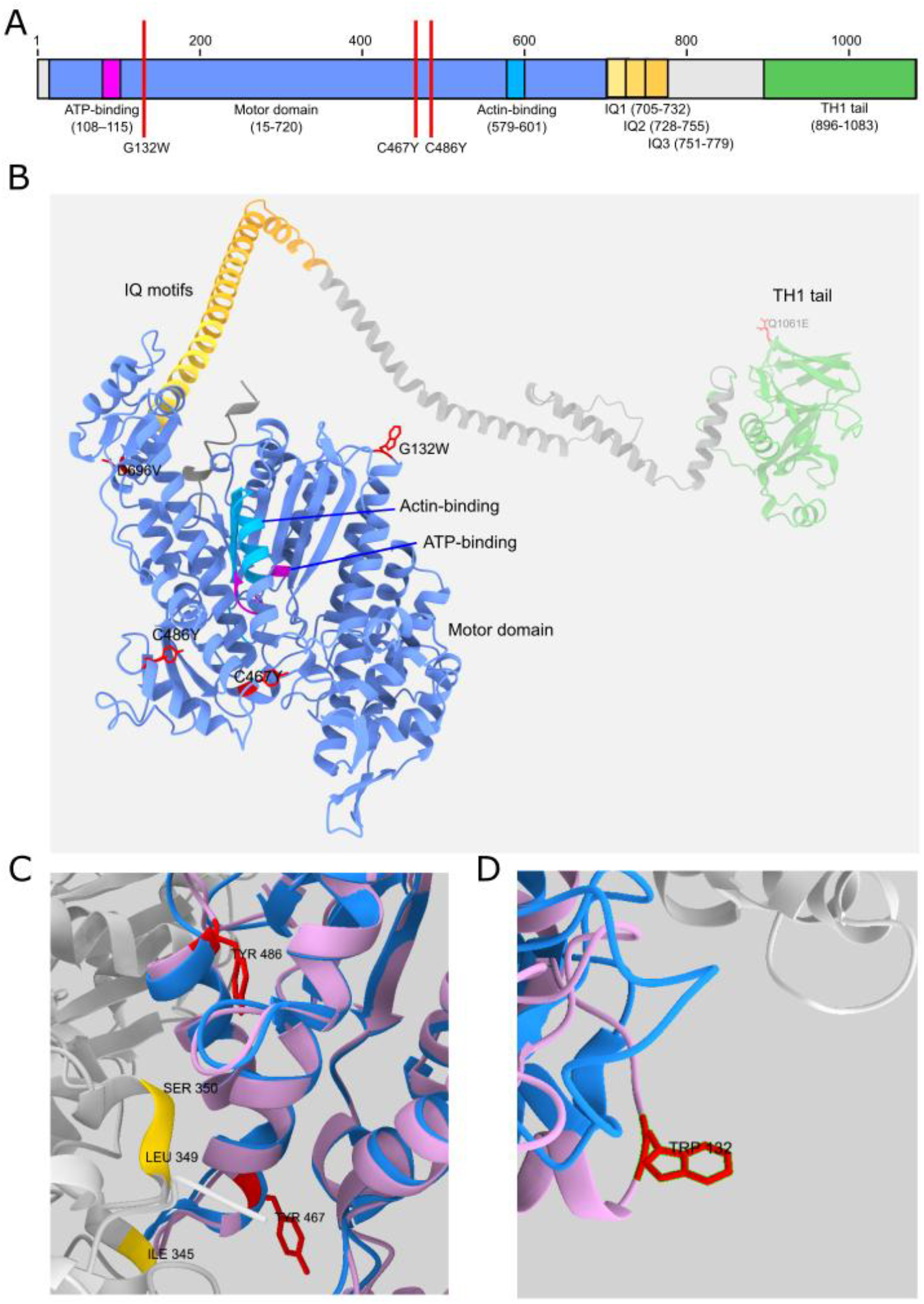
Functional annotation and modelling of the myosin I a/b gene in *Euhadra*. **(A)** The predicted domain structure. **(B)** 3D model highlighting non-conservative amino acid substitutions, three of which are found in the motor domain in all sinistral *Euhadra* species (G132W, C467Y, C486Y), and two of which are found in a subset of *Euhadra* species, including *E. quaesita* (Q1061E, D696V). **(C)** and **(D)** sinistral and dextral myosins aligned to structure of actin-bound myosin, showing key amino acid changes. Sinistral myosin is in pink, dextral in blue. In **(C)** grey indicates myosin, showing key positions. In **(D)** grey indicates calmodulin.

The sinistral version of the myosin in *Euhadra* has multiple amino acid differences compared with other *Euhadra* (**Figure 3C**). Specifically, the sinistral allele is defined by eight amino acid changes that are fixed in all sinistral isolates and species of *Euhadra*, and not found in any dextral *Euhadra*. To further explore these coding differences, structural modelling was used to identify changes that are likely functionally important. Of the amino acid changes that are only found in the sinistral lineage (**Table S10**), three are non-conservative (G132W, C467Y, C486Y). An analysis using Provean^28^ showed that the same three changes are likely functionally important (G132W -5.026; C467Y -10.392; C486Y -2.650). Glycine (G) is very small and flexible, whereas tryptophan (W) is large and aromatic. Cysteine (C) has a thiol that can form disulfide bonds and is small, whereas tyrosine (Y) is a bulky aromatic with a polar OH.

In comparison to the eight amino acid chances that define the sinistral group, neither the dextral monophyletic group that comprises *E. brandtii, E. senckenbergiana, E. herklotsi, E. dixoni*, and *E. awaenesis*, or the dextral lineage of *E. quaesita*, has any lineage-specific amino acid changes (**Figure 3C**). Reflecting these differences, there is strong evidence for a relaxation of selection in the sinistral lineage. Using RELAX, we detected a pronounced relaxation of selective constraint along the focal branches (K = 0.24, LR = 11.63, p = 0.001), whereas background branches were dominated by strongly purifying selection (ω ≈ 0.05– 0.07), the focal branches showed a substantial upward shift in ω values (ω ≈ 0.49), consistent with a genome-wide weakening of purifying selection rather than adaptive evolution. A complementary BUSTED analysis, incorporating site-to-site synonymous rate variation, found no evidence for gene-wide episodic diversifying selection (*P* = 0.237). Allowing ω > 1 did not significantly improve model fit, indicating that the observed elevation in ω reflects a weakening of purifying selection rather than adaptive evolution. Together, these analyses provide strong evidence that the sinistral lineage has experienced relaxed selective constraint without detectable episodic positive selection.

Superimposition of sinistral and dextral myosin models showed identical backbones and thus unchanged global architecture (RMSD 0.058 Å over 1083 pairs). At the eight variant sites, no steric clashes were detected and Cα-Cα separations were all < 0.35 Å, indicating no gross conformational change. In comparison, models indicate that the same three substitutions in the motor domain (G132W, C467Y, C486Y; **Figure 4B**) may reshape local hydrogen-bonding networks, indicating changes in stiffness, compliance or coupling efficiency (**Table S10**).

To further explore the putative function of the three amino-acid substitutions, the sinistral and dextral models were mapped onto a structure of myosin-1c bound to actin in the ADP-bound state^29^ (PDB ID: 9CFU). To assess whether sinistral- and dextral-specific substitutions alter the global motor-domain structure, both models were first aligned to the actin-bound myosin-1c cryo-EM structure motor domain, using core residues (15–700) and also locally (457-477, 476-496, 122-142). Across the whole alignment pruned RMSDs were approximately 1.06–1.07 Å, indicating near-identical backbone conformations. Thus, neither variant exhibits large-scale structural divergence relative to the experimental structure. Similarly, there is no evidence that residue 467 (RMSD 0.38/0.40 Å in dex/sin), 486 (0.55/0.75 Å) or 132 (1.26/1.24 Å) induce any local backbone rearrangement in sinistral or dextral myosin relative to the myosin-1C structure.

In the sinistral model, residue 486 occupies an interaction-permissive spatial environment (∼7 Å) proximal to actin residue Ser350, whereas there is much more distant proximity in the dextral variant (∼10 Å; **Figure 4C**). This geometry suggests a potential for conditional or transient interactions. Residue 467 also lies on the canonical actin-binding surface of the motor domain and occupies a position proximal to actin subdomain 1 in actin-bound myosin structures, with the sinistral version potentially closer to actin residues Leu349 and Ile345 (**Figure 4C**). Finally, the G132W substitution introduces substantial steric bulk at an outward-facing position proximal to the motor-lever junction (**Figure 4D**). Given the known flexibility and strain sensitivity of myosin-I lever arms, this mutation may influence mechanical coupling between the motor domain and the calmodulin-stabilised lever, even in the absence of a fixed interaction in the static structure. Together, these analyses demonstrate that sinistral-specific substitutions cluster within functional interaction regions of the actomyosin complex, altering local interface geometry rather than global structure.

As a point of comparison, positions 132 and 467 are invariable in 57 available gastropod transcriptomes, with position 486 also invariable, except in one species (C486I; *Onchidoris muricata*). Similarly, myosin I a/b sequences from sinistral and dextral Hygrophila snails (*L. stagnalis* versus *Biomphalaria glabrata*) showed 66 fixed amino acid changes between them, but all are predicted to be neutral by Provean. Finally, the region around position 467 is the conserved; all species from across the Bilateria invariably have a cytosine at that position.

### Molluscan LR asymmetry in context

Left-right asymmetry in animals ultimately originates from molecular and cellular chirality, yet the earliest symmetry-breaking events remain poorly understood. Previous work has identified formin as the causative gene for pathological change in *L. stagnalis*, implicating actin-based cytoskeletal processes in determining the handedness of early cleavage, which is consistent with wider studies.^3^ These discoveries have supported a broader model in which LR asymmetry arises from inherently chiral cytoskeletal dynamics, particularly involving actin and myosin.^6,30^ The discovery of a myosin I a/b isoform as causative of variation in the LR asymmetry of the land snail *E. quaesita* represents a major advance in understanding the mechanistic diversity of LR asymmetry for two reasons. First, although unconventional myosins have been implicated in LR patterning across invertebrate and vertebrate taxa^31,32^, notably *Drosophila* and zebrafish^10,33-35^, no previous study has implicated a class I a/b myosin in directing LR asymmetry in any animal^26^. Second, our results show that, despite acting within the same broad cytoskeletal pathway as formin, snail chiral variation can be specified by fundamentally different molecular components across lineages. The finding therefore broadens the known repertoire of symmetry-breaking factors and illustrates how distinct cytoskeletal proteins can converge on the same developmental outcome.

These findings provide a framework for understanding how cytoskeletal asymmetry is translated into organismal form, and they illustrate the potential for applying similar approaches to the many gastropod lineages that vary in chirality but have not yet been genetically characterised.

An important next step is to determine the functional consequences of the snail sinistral myosin I variant. To gain final proof of function of the individual amino acid changes, it would be ideal to engineer a snail with the individual amino acid changes. Unfortunately, no study in snails has manipulated coding variation, with only a few CRISPR-Cas9 papers in eleven years, all of which are knockdowns. In any case, since a major aim is to understand the generality of these findings, an alternative strategy should be to use more powerful gene-editing methods in other species to identify the single amino acid that changes chirality, and whether permissive or background changes are a necessary precursor. In *Drosophila*, the direction of organ twisting depends on the biochemical and biophysical properties of specific myosin I paralogs: Myo 1D drives left-handed actin rotation *in vitro*, whereas Myo 1C favours the opposite direction.^10^ ATPase function differs between the different forms.^36^ In other experiments, it has been shown that myosins with chiral activity can autonomously organize actin filaments into stable, chiral structures through collective motion.^9^ If the *Euhadra* myosin variant similarly alters motor activity, actin-myosin coupling, or the direction of filament rotation, this could provide a direct mechanistic explanation for reversed cleavage chirality.

The findings raise the question of what would happen if similar mutations were introduced into *Caenorhabditis elegans*, in which chirality is due to an early spiral twist^2^, *Drosophila*, or vertebrate systems? Would they generate mirror-image LR asymmetry, as in the sinistral nematode^37^, or would developmental buffering suppress such effects? On the one hand, given that myosins and formins are deeply conserved, and functionally related, it is conceivable that altering the chirality of actin-based dynamics could induce reversed morphogenesis in other taxa. Alternatively, developmental context and tissue architecture may constrain such outcomes. Testing these ideas experimentally will provide important insight into the evolutionary plasticity of symmetry-breaking systems.

Together, our results show that a previously unrecognised member of the unconventional myosin I family underlies natural inherited variation in LR asymmetry in *E. quaesita*, expanding the known diversity of cytoskeletal mechanisms capable of establishing organismal chirality. This discovery deepens our understanding of how molecular-scale asymmetries propagate to the level of the whole body and highlights snails as an exceptional model for dissecting the fundamental origins of left and right.

## Data and code availability

The genome assembly (GCA_982298635) and raw sequence reads are available via bioproject PRJEB106571 and sample ERS28485073. Raw whole genome sequence reads of other individuals are available via bioproject PRJEB110179. Scripts will be available at https://github.com/AlecLewis-Bio/EuhadraGenomics.

## Acknowledgements

This work was supported by the University of Nottingham, the Biotechnology and Biological Sciences Research Council, via a studentship to MJ, grant number BB/M008770/1, and a studentship to AL, grant number BB/T0083690/1. Collaborative work was enabled by a Daiwa Foundation small grant to AD (6495/15181), a JSPS/British Council Summer studentship to AL. The work was also enabled by a prior BBSRC studentship to Paul Richards, with additional funding from BBSRC grant BB/F018940/1, the Japanese Society for the Promotion of Science / Royal Society 2+2 award, and the Genetics Society to AD. Bioinformatic analyses were enabled by a UoN-funded MSc in Bioinformatics (to AD), and access to the University of Nottingham HPC.

## Author contributions

Conceptualisation: AD, SC. Field work: AD, AL, YI, SI, KK, SC, TH. Laboratory work: AD, AL, YI, MJ, CM, TH, DJJ. Bioinformatics: AD, AL, YI. Data curation: AD, YI. Writing – original draft: AD. Writing – review and editing: AD, AL, YI, AH, CM. Visualization: AD, YI, AL. Supervision: AD, SC. Funding acquisition: AD, SC, YI, TH.

## Supplementary Methods

Detailed methods are provided in the online version of this paper.

## References

1. Naganathan, S.R., Popovic, M., and Oates, A.C. (2022). Left-right symmetry of zebrafish embryos requires somite surface tension. Nature 605, 516-+. 10.1038/s41586-022-04646-9.

2. Khor, M., Rain, Y., Sangha, G., and Sugioka, K. (2025). Cadherin flow translates cortical chirality into left-right asymmetric cell division dynamics in Caenorhabditis elegans embryos. Current Biology 35, 5320–5331. 10.1016/j.cub.2025.09.052.

3. Tsikolia, N., Nguyen, D.T.L., and Tee, Y.H. (2025). Mechanisms of left–right symmetry breaking across scales. Current Opinion in Cell Biology 95, 102564. 10.1016/j.ceb.2025.102564.

4. Katoh, T.A., Omori, T., Mizuno, K., Sai, X., Minegishi, K., Ikawa, Y., Nishimura, H., Itabashi, T., Kajikawa, E., Hiver, S., et al. (2023). Immotile cilia mechanically sense the direction of fluid flow for left-right determination. Science 379, 66–71. 10.1126/science.abq8148.

5. Djenoune, L., Mahamdeh, M., Truong, T.V., Nguyen, C.T., Fraser, S.E., Brueckner, M., Howard, J., and Yuan, S. (2023). Cilia function as calcium-mediated mechanosensors that instruct left-right asymmetry. Science 379, 71–78. 10.1126/science.abq7317.

6. Tee, Y.H., Goh, W.J., Yong, X.B., Ong, H.T., Hu, J.R., Tay, I.Y.Y., Shi, S.D., Jalal, S., Barnett, S.F.H., Kanchanawong, P., et al. (2023). Actin polymerisation and crosslinking drive left-right asymmetry in single cell and cell collectives. Nature Communications 14, 776. 10.1038/s41467-023-35918-1.

7. Tee, Y.H., Shemesh, T., Thiagarajan, V., Hariadi, R.F., Anderson, K.L., Page, C., Volkmann, N., Hanein, D., Sivaramakrishnan, S., Kozlov, M.M., and Bershadsky, A.D. (2015). Cellular chirality arising from the self-organization of the actin cytoskeleton. Nature Cell Biology 17, 445–457.

8. Inaki, M., Liu, J., and Matsuno, K. (2016). Cell chirality: its origin and roles in left– right asymmetric development. Philos Trans R Soc Lond B Biol Sci 371, 2015.0403. 10.1098/rstb.2015.0403.

9. Haraguchi, T., Yoshimura, K., Inoue, Y., Imi, T., Hasegawa, K., Nagai, T., Furusawa, H., Mori, T., Matsuno, K., and Ito, K. (2026). Elucidating chiral myosin-induced actin dynamics: From single-filament behavior to collective structures. Proceedings of the National Academy of Sciences of the United States of America 123, e2508686123. 10.1073/pnas.2508686123.

10. Lebreton, G., Geminard, C., Lapraz, F., Pyrpassopoulos, S., Cerezo, D., Speder, P., Ostap, E.M., and Noselli, S. (2018). Molecular to organismal chirality is induced by the conserved myosin 1D. Science 362, 949–952. 10.1126/science.aat8642.

11. Da Silva Anjos Machado, X., Francis, R.M., Mandhar, K., Perinparajah, S., Yadati, S.S., and Brand, T. (2025). The Role of Left-Right Asymmetry in Congenital Heart Defects: Taking the Right Turn. Curr. Treat. Options Cardiovasc. Med. 27, 57. 10.1007/s11936-025-01114-1.

12. Davison, A. (2020). Flipping shells: unwinding LR asymmetry in mirror-image molluscs. Trends in Genetics 36, 189–202. 10.1016/j.tig.2019.12.003.

13. Sturtevant, A.H. (1923). Inheritance of direction of coiling in Limnaea. Science 58, 269–270.

14. Boycott, A.E., and Diver, C. (1923). On the inheritance of sinistrality in Limnaea peregra. Proc R Soc Biol Sci Ser B 95, 207–213.

15. Martín-Durán, J.M., and Marlétaz, F. (2020). Unravelling spiral cleavage. Development 147, dev181081. 10.1242/dev.181081.

16. Davison, A., McDowell, G.S., Holden, J.M., Johnson, H.F., Koutsovoulos, G.D., Liu, M.M., Hulpiau, P., Van Roy, F., Wade, C.M., Banerjee, R., et al. (2016). Formin is associated with left-right asymmetry in the pond snail and the frog. Current Biology 26, 654–660. 10.1016/j.cub.2015.12.071.

17. Abe, M., and Kuroda, R. (2019). The development of CRISPR for a mollusc establishes the formin Lsdia1 as the long-sought gene for snail dextral/sinistral coiling. Development 146, dev.175976. 10.1242/dev.175976.

18. Noda, T., Satoh, N., and Asami, T. (2019). Heterochirality results from reduction of maternal diaph expression in a terrestrial pulmonate snail. Zoological Letters 5, 2, 2. 10.1186/s40851-018-0120-0.

19. Davison, A., Barton, N.H., and Clarke, B. (2009). The effect of coil phenotypes and genotypes on the fecundity and viability of Partula suturalis and Lymnaea stagnalis. J Evol Biol 22, 1624–1635. 10.1111/j.1420-9101.2009.01770.x.

20. Utsuno, H., Asami, T., Van Dooren, T.J.M., and Gittenberger, E. (2011). Internal selection against the evolution of left-right reversal. Evolution 65, 2399–2411. 10.1111/j.1558-5646.2011.01293.x.

21. Gittenberger, E., Hamann, T.D., and Asami, T. (2012). Chiral speciation in terrestrial pulmonate snails. PLoS One 7, e34005, e34005. 10.1371/journal.pone.0034005.

22. Davison, A., Chiba, S., Barton, N.H., and Clarke, B.C. (2005). Speciation and gene flow between snails of opposite chirality. Public Library of Science Biology 3, e282.

23. Ueshima, R., and Asami, T. (2003). Single-gene speciation by left-right reversal - A land-snail species of polyphyletic origin results from chirality constraints on mating. Nature 425, 679–679.

24. Davison, A., Barton, N.H., and Clarke, B. (2009). The effect of coil phenotypes and genotypes on the fecundity and viability of Partula suturalis and Lymnaea stagnalis: implications for the evolution of sinistral snails. Journal of Evolutionary Biology 22, 1624–1635. 10.1111/j.1420-9101.2009.01770.x.

25. Hoso, M., Kameda, Y., Wu, S.-P., Asami, T., Kato, M., and Hori, M. (2010). A speciation gene for left-right reversal in snails results in anti-predator adaptation. Nature Communications 1, ncomms1133, 133. 10.1038/ncomms1133.

26. Okumura, T., Sasamura, T., Inatomi, M., Hozumi, S., Nakamura, M., Hatori, R., Taniguchi, K., Nakazawa, N., Suzuki, E., Maeda, R., et al. (2015). Class I Myosins Have Overlapping and Specialized Functions in Left-Right Asymmetric Development in Drosophila. Genetics 199, 1183–U1493. 10.1534/genetics.115.174698.

27. Utsunomiya, S., Takebayashi, K., Yamaguchi, A., Sasamura, T., Inaki, M., Ueda, M., and Matsuno, K. (2024). Left-right Myosin-Is, Myosin1C, and Myosin1D exhibit distinct single molecule behaviors on the plasma membrane of Drosophila macrophages. Genes Cells 29, 380–396. 10.1111/gtc.13110.

28. Choi, Y., and Chan, A.P. (2015). PROVEAN web server: a tool to predict the functional effect of amino acid substitutions and indels. Bioinformatics 31, 2745–2747. 10.1093/bioinformatics/btv195.

29. Chavali, S.S., Carman, P.J., Shuman, H., Ostap, E.M., and Sindelar, C.V. (2025). High-resolution structures of Myosin-IC reveal a unique actin-binding orientation, ADP release pathway, and power stroke trajectory. Proceedings of the National Academy of Sciences 122, e2415457122. doi:10.1073/pnas.2415457122.

30. Naganathan, S.R., Fürthauer, S., Rodriguez, J., Fievet, B.T., Jülicher, F., Ahringer, J., Cannistraci, C.V., and Grill, S.W. (2018). Morphogenetic degeneracies in the actomyosin cortex. eLife 7, e37677. 10.7554/eLife.37677.

31. Tingler, M., Kurz, S., Maerker, M., Ott, T., Fuhl, F., Schweickert, A., LeBlanc-Straceski, J.M., Noselli, S., and Blum, M. (2018). A conserved role of the unconventional myosin 1d in laterality determination. Current Biology 28, 810-816.e813. 10.1016/j.cub.2018.01.075.

32. Yuan, S., and Brueckner, M. (2018). Left-right asymmetry: myosin 1D at the center. Current Biology 28, R567–R569. 10.1016/j.cub.2018.03.019.

33. Hozumi, S., Maeda, R., Taniguchi, K., Kanai, M., Shirakabe, S., Sasamura, T., Speder, P., Noselli, S., Aigaki, T., Murakami, R., and Matsuno, K. (2006). An unconventional myosin in Drosophila reverses the default handedness in visceral organs. Nature 440, 798–802.

34. Spéder, P., Ádám, G., and Noselli, S. (2006). Type ID unconventional myosin controls left–right asymmetry in Drosophila. Nature 440, 803–807. 10.1038/nature04623.

35. Juan, T., Geminard, C., Coutelis, J.B., Cerezo, D., Poles, S., Noselli, S., and Furthauer, M. (2018). Myosin1D is an evolutionarily conserved regulator of animal left-right asymmetry. Nature Communications 9, 1942. 10.1038/s41467-018-04284-8.

36. Báez-Cruz, F.A., and Ostap, E.M. (2023). Drosophila class-I myosins that can impact left-right asymmetry have distinct ATPase kinetics. J Biol Chem 299, 104961. 10.1016/j.jbc.2023.104961.

37. Felix, M.A., Sternberg, P.W., and Ley, P.d. (1996). Sinistral nematode population. Nature 381, 122.

