## Supplementary Information for "An ancient polymorphism in myosin I a/b determines the left-right asymmetry of Japanese snails"

### Methods

#### Samples

*Euhadra* is genus of around 30 species of snails, for which four species are always sinistral (*E. decorata*, *E. grata*, *E. murayamai*, *E. scaevola*) and one species, *E. quaesita*, shows chiral variation, with the dextral form sometimes called *E. aomoriensis*.[1] Like other land snails, *Euhadra* are simultaneous hermaphrodites and usually mate reciprocally, as male and female at the same time.

In previous work, we identified two regions in northern Japan (Iwate and Yamagata) where both chiral types of *E. quaesita* are physically close together and may come into contact, with some evidence of recent gene-flow between them.[2] Here, we identified two further possible contact zones between dextral and sinistral snails, in Fukushima and Niigata prefectures, making four total (**Figure 1D**). The last discovered site (#4) in Niigata is key to identifying the causal gene because (unlike the others) we found a broad zone of sympatry where both chiral types are found together (**inset to Figure 1D)**, also having shells that look outwardly similar (**Figure 1A**).

The main dataset was derived from snails from these four sites, plus a collection of multiple species of *Euhadra*, including all sinistral species. These were all sequenced to 5-10 x depth using Illumina sequencing (**Table S2**). A second dataset was an independent and larger set of *E. quaesita* to which double digest restriction site-associated DNA sequencing (ddRAD) methods were applied (**Table S11**).

#### The culture of snails

*Euhadra* snails were kept under standard conditions in the laboratory, including *ad libitum* fresh carrot or sweet potato and a dry powder mix made up of sow weaner pellets, milk powder, chalk and nutritional yeast in a 3:3:3:1 ratio. To record mating, snails were isolated for some weeks and then placed in an arena; full details will be published separately. To collect egg batches and derive single cell embryos, individual *Euhadra* snails were isolated and observed for digging behaviour. Upon finding a laying individual, whole batches of eggs were removed, counted and then stored separately until they hatched, approximately one month later, whereupon chirality phenotype and percentage hatch of all offspring was recorded. To recover single cell embryos, freshly laid eggs were removed. The egg capsule was broken open with fine forceps and the single cell embryo removed with a P2 pipette. Capsular fluid that tends to adhere to the embryo was removed by pipetting the embryo in sterile water, a necessary step to prevent interference with downstream molecular biological methods. The embryo was then placed in 2µl of PBS and frozen at -70°C for subsequent use.

For comparison, we also derived single cell embryos for sinistral and dextral pond snails *Lymnaea stagnalis*, using the same method as described previously.[3] A key difference compared with *Euhadra* is that the pond snails are from a dextral laboratory line into which the sinistral allele was introgressed over many generations. The two lines are therefore mostly genetically homogenous, except at the chirality locus.

#### Inheritance of chirality in *Euhadra* and tests for pathology

In all previous studies, chirality in snails is determined by locus of maternal effect, with either dextral or sinistral dominant.[4] In *Euhadra* the dominance is not known. The difficulty in understanding dominance is that as populations are fixed for their chirality, interchiral laboratory mating of virgins is a required first step. Subsequently, several generations of breeding are then required to establish dominance.

As a starting point to understanding chirality in *Euhadra*, *E. quaesita* individuals from the newly discovered contact zone (#4) were returned to the laboratory and allowed to lay eggs. Following the convention, we recorded and described maternal phenotype as either dextral or sinistral, and the inferred genotype, based upon the chirality of the offspring, in letters (*D* or *S*). If dextral is dominant, then the five phenotype/genotype combinations are dex*DD* dex*DS* dex*SS* and sin*SS* sin*DS*; a "sin*DD*" snail is not possible because the mother would have to have had the dominant *D* allele. Similarly, if sinistral is dominant, then the five phenotype/genotype combinations are dex*DD* dex*DS* and sin*SS* sin*DS* sin*DD* (no dex*SS*).

In other species, gross pathology of the sinistral mutation is evident because a high proportion of the eggs do not hatch.[5,6] Therefore, to confirm that the mutation is not pathological in *E. quaesita*, clutches of eggs were counted and monitored for both dextral and sinistral snails from the Niigata sympatric site and the Fukushima allopatric sites. The maternal genotype was recorded, upon egg hatching and the number of viable individuals counted. Differences in average hatch rate between chiral types and sampling location were assessed with Mann-Whitney U tests. Additionally, we used a general linear mixed model with beta-regression, either including or not-including an interaction factor between chirality and sample location.

#### *De novo* genome assembly and annotation

The genome assemblies and annotation will be described in detail in a separate publication. In brief, high molecular weight DNA was extracted from the foot tissue of an individual *E. quaesita* from Kumagane, Sendai, Miyagi (38.29359942°N 140.6890908°E) using the QIAGEN Genomic-tip 100/G kit. A PacBio library was prepared via service provider Novogene, and then Revio HiFi reads derived via three SMRT Cells. A limited number of long reads were also derived from another individual using an Oxford Nanopore MinION. To scaffold the reads, a Hi-C library was prepared from the DNA of an offspring of the PacBio snail, using the Proximo Hi-C kit (Animal), and then sequenced using standard Illumina methods.

*De novo* assembly was initially performed with Hifiasm 0.19.7-r588[7] using HiFi and Nanopore reads. Haplotigs were purged by purge_dups 1.2.5[8], applying the "-e" option in the get_seqs module. The Hifiasm assembly was then used as input for Hi-C scaffolding. Hi-C reads were mapped onto the assembly using bwa 0.7.17-r1188[9] and samtools 1.18[10]. Then, the mapped reads were used for Hi-C scaffolding using YaHS 1.2.2[11]. The contact map was visualized, and one mis-joined scaffold was edited. The final chromosome-scale scaffolding was performed using the HapHiC v1.0.1 pipeline[12], which includes the bundled Juicer version packaged with HapHiC v1.0.1 for Hi-C–based correction and scaffolding. The resulting scaffolds were further processed using Juicer 1.2[13], Juicer Tools 1.9.9, and visualised and manually curated in Juicebox 2.15.0.0.[14] These steps together produced the final edited assembly used for downstream analyses.

Following genome assembly, RepeatModeler 2.0.[15] was used to build a species-specific repeat content library, and the repeat content was masked using RepeatMasker 4.2.3. Both software tools were implemented in TETools 1.96[16]. Gene prediction was performed for the soft-masked sequence using BRAKER 3.0.8[17]. Available RNA-seq data (NCBI accession numbers SRR) was used with new RNA-seq data derived from single cell embryos (see below), and protein data, molluscan proteomes in OrthoDB 11[18]) to annotate the genome. The gene content completeness of the assemblies and the transcriptome was assessed using BUSCO 5.7.1[19], by searching against the metazoan core genes (metazoa_odb10).

#### Whole genome re-sequencing, mapping to reference, variant calling and filtering

The genome of individual snails was sequenced by service provider Novogene, using Illumina paired-end methodology (NovaSeq 6000 PE150), and aiming for ~5-10x depth. FASTP v0.23.2 software was used in pre-processing of fastq files, specifically to check the quality, and for adapter trimming and quality filtering of the reads.[20] Then, the read files were aligned to the *E. quaesita* genome, using the Burrows-Wheeler Alignment method in bwa 0.7.17-r1188[9], with default settings and marking shorter split hits as secondary (-M). Samtools 1.18[10] was then used to sort and compress the sam files to bam files. Picard tools 3.0-Java-17[21] was used to remove putative duplicate reads in each bam file, then bcftools v1.18[10] was used to call variants and create a vcf file. To filter the vcf files, vcftools 0.1.16[22] was used, removing indels, retaining only biallelic sites with MAF ≥ 0.05, QUAL ≥ 30, and mean depth between 3 and 10; individual genotypes outside the depth range 3-10 were set to missing. Various datasets were then output, by varying the required proportion of non-missing genotypes. 85% was used for most analyses, because it is a good compromise between number of SNPs and degree of missingness.

#### Genome-wide association methods

Prior to conducting genome-wide association analyses individual snails were scored for their chirality phenotype and genotype. One of the difficulties in doing this is that the maternal phenotype (dextral or sinistral coiling) does not always correspond to the individual genotype (*D* or *S*), as inferred by the direction of coiling of the hatched offspring from a mother [23], because of the maternal inheritance.

In species and locations where chirality is fixed then the assumption is that the chirality phenotype represents the chirality genotype, because otherwise reversed chirality individuals would sometimes be present. Thus, where chirality is fixed, all dextral snails are genotype *DD* and all sinistral snails are genotype *SS*.

In comparison, in regions of gene-flow between sinistrals and dextrals, the phenotype may not correspond to the chiral genotype. The first strategy to score genotype was to infer it from the offspring chirality; a dextral snail that produces sinistral offspring must have at least one *S* allele, and *vice versa*, irrespective of dominance relations. Previously, it has been shown using a mathematical model and also empirical data that phenotype is a predictor of genotype, even when there is geneflow[5,23]. Therefore, in the absence of offspring, the second strategy was to score the genotype as the same as the phenotype, knowing that this would come with some error. This dataset of *D* or *S* snails was then used for the genome wide association analyses GWAS ("Maternal genotype" in Table 2).

Subsequently, we were able to infer the full chirality genotype of snails that underwent whole genome sequencing, based on the chromosome 27 haplotypes ("Inferred chirality genotype" in Table 2; see Figure 3A for explanation). This information was of course not used for the GWAS.

A genome-wide association method was performed in R using the Gaston 1.6 package[24], applied to the main dataset of SNPs, including multiple individuals from the #4 Niigata contact zone. Gene-flow means that recombination events that have occurred over generations of ongoing gene-flow should enable association mapping methods, with the size of region depending upon the number of generations of introgression and local recombination rate.

In brief, the binary genotype records were imported into Gaston, alongside the VCF file. Prior to association testing, genotypes were standardised, and a genetic relationship matrix

(GRM) was estimated to account for population structure. Association tests were then conducted using Gaston's linear mixed model (LMM) framework with a binary response, incorporating the GRM as a random effect. Genome-wide significance was evaluated using a strict Bonferroni-adjusted threshold, based on the number of SNPs tested, as well as a less conservative FDR-adjusted method, using the Benjamini–Hochberg (BH) correction. Manhattan plots were generated directly from Gaston output, both genome-wide and for specific chromosomes or chromosomal subregions of interest. Significant SNPs were extracted by filtering for *P* values below the multiple-testing-adjusted threshold, and regional plots were produced by sub-setting association results to defined chromosomal intervals. To assess the distribution of test statistics and potential inflation, quantile-quantile (QQ) plots were generated using the same association results. SNPs surpassing the multiple-testing-corrected threshold were extracted for further investigation.

#### Evolutionary genomics

Evolutionary-genomic methods and statistics were used to further corroborate regions of the genome differentiated between dextral and sinistral individuals.

First, a sliding-window analysis was first performed across the genome to calculate Fst and Dxy, allowing detection of windows showing elevated differentiation or divergence between the two phenotypic groups. For this, the vcfs were first split into each of the 28 chromosomes using bcftools (v.1.18), and subsetted to include only *E. quaesita*[10]. The files were then filtered with vcftools (v.0.1.16), applying the same quality and depth filters as used for the GWAS, and allowing for up to 5% missing data, but retaining invariant sites. A minor allele frequency was not applied as invariant sites were needed for calculations of pi ($\pi$). Haplotype phasing was carried out using beagle (v.5.5, impute = true, window size = 50kb, overlap = 5kb), with data converted from the vcf file format to Simon Martin's genotype file format (.geno)[25]. Genome wide patterns of genomic differentiation (Fst) and absolute divergence (Dxy) were calculated between all populations of *E. quaesita* using the popgenWindows.py (window = 50kb, step = 50kb, minimum sites = 2500) script, available at https://github.com/simonhmartin/genomics_general. We applied 99th and 99.9th percentile cutoffs as markers of significance, justifying the high thresholds due to chirality being a single Mendelian locus.

Second, for phylogenomic analyses, we used the same filtered vcf files, allowing for up to 5% missing data, and using a minor allele frequency (0.03) and quality score minimum of 30), converting the vcf to Phylip format using the vcf2phylip.py script[26]. For each alignment, IQ-TREE2[27] was run with automated model selection and ultrafast bootstrap approximation to assess node support. Phylogenies were inferred under the best-fitting substitution model identified by the software, with ultrafast bootstrap support calculated across 1000 replicates. Whole-genome trees were used to reconstruct overall relationships among samples, whereas region-restricted trees allowed comparison of local genealogical structure across the genome. One limitation of a concatenated phylogeny is that it assumes a single underlying topology. Therefore, an alternative approach was taken to generate a species tree using methods consistent under the multispecies coalescent theory. For this, each chromosome filtered vcf was subset into windows spanning 500 kb. The individual windows were converted to Phylip format, and maximum likelihood phylogenies were generated using IQ-TREE2, using the modelfinder option (-m MFP) to find the optimum nucleotide substitution model for each window individually. The best trees found by IQ-TREE2 for each window were concatenated into a file to be used as input for ASTRAL (v.5.7.8)[28]. ASTRAL was then used to generate a final species tree based on the varying phylogenetic topologies seen in windows across the genome, support is given in local posterior probability.

Third, a topology weighting analysis was used to quantify heterogeneity in genealogical relationships across the genome and to explicitly compare support for alternative evolutionary histories that are obscured in a single genome-wide tree[25,29]. The intention was to identify if there are genomic regions where samples cluster by chirality rather than by geography, contrary to the genome-wide species tree. Using the same haplotype data, TWISST was applied to a collection of phylogenetic trees generated with phyml_sliding_windows.py for non-overlapping windows containing 50 variable sites. Specifically, we used a four-population model (Yamagata sinistrals, Yamagata dextrals, Fukushima dextrals, Fukushima sinistrals) to test whether local genealogies supported the expected geography-based species tree or instead supported alternative topologies grouping individuals by chirality across regions. Genomic regions with a topology weight of 1 in support of chirality-based groupings were interpreted as loci whose evolutionary history deviates from the genome-wide pattern and may reflect trait-associated introgression or shared ancestry linked to shell chirality.

We also undertook several other analyses to better understand the genomic context and also historical phylogeography of the snails.

Linkage disequilibrium was measured ($r^2$) between sites using Plink (v.1.9)[30] for pairs of SNPs that were separated by a distance ranging from 100 basepairs to 1 megabase. The calculation was run using on *E. quaesita* samples and run individually for each chromosome (subsetted during analysis with the Plink –chr function). Average linkage disequilibrium for given intervals across the genome was calculated using a custom python script, available from: https://github.com/speciationgenomics/scripts/blob/master/ld_decay_calc.py.

The same data set was also used to explore admixture between chirally variable *E. quaesita* sample sites. First, Plink (v.1.9) was used to prune the data for linkage disequilibrium across all chromosomes, filtering in 50 kb windows for $r^2$ values over 0.1. Admixture proportions were calculated in ADMIXTURE (v.1.3.0) for up to 10 ancestral populations (K) [31], using the .bed file as input. Cross-validation (CV) error, as calculated in ADMIXTURE was used to identify the optimal value of K. Additionally, a PCA was run using the --pca option in Plink, as a non-model based method to determine whether snails group by geography or chirality.

#### Double digest restriction site-associated DNA sequencing

A set of *E. quaesita* individuals (n=107; Table 11) were put through double digest restriction site-associated DNA sequencing, or ddRAD-seq[32], using the same method as previous[33]. In brief, libraries were prepared following an established protocol[34], with slight modifications, using the restriction enzymes EcoRI and MspI to digest total DNA. Libraries were sequenced for 150 bp paired-end reads using a DNBSEQ-G400 by BGI (China). Demultiplexing, quality control and *de novo* assembly used ipyrad 0.9.108 [35]. Quality control then used trimmomatic 0.39, with the options "ILLUMINACLIP:NexteraPE-PE.fa:2:30:10 LEADING:30 TRAILING:30 MINLEN:140 CROP:140". Read-mapping used bwm-mem2 2.2.1, with stacks 2.62 used to assemble[36]. Loci shared by 70% of 130 individuals were then output as genotypes.

#### Single cell RNA-seq methods

Individual single cell embryos were processed for full-length single-cell transcriptome profiling using the SMART-Seq3 protocol[37]. Briefly, first-strand cDNA synthesis and incorporation of unique molecular identifiers (UMIs) were performed according to the SMART-Seq3 chemistry, enabling capture of both 5′ UMI-tagged fragments and full-length cDNA. Following reverse transcription, cDNA was amplified by PCR, and each cell was processed as an independent library. Libraries were prepared using standard tagmentation-based workflows, with sample identity encoded by unique i7/i5 index combinations. Because SMART-Seq3 uses index barcodes rather than read-embedded barcodes, cell assignment was performed based on index pairs rather than extraction from R1/R2 read sequences. Final libraries were pooled in equimolar amounts and sequenced in paired-end mode on an Illumina platform to a depth of approximately 0.5-2 million reads per cell.

Reads were demultiplexed by index combinations. Then, raw sequencing reads were processed with Fastp to perform adapter trimming, quality filtering, and removal of low-quality bases[20]. Reads passing quality control were then filtered using SortMeRNA [38] to remove ribosomal RNA and other high-abundance sequences. The remaining reads were then aligned to the reference genome using HISAT2 2.2.1[39], using a maximum allowed intron length of 60kb, and taking a conservative approach in not allowing discordant paired-end alignments or cases where only one mate aligns.

Aligned reads for each individual embryo were assembled into transcript models using StringTie 2.2.1.[40] Individual sample GTF files were merged with the StringTie merge function to generate a combined transcriptome annotation, from which a custom transcriptome reference was derived. Gene-level quantification was performed using featureCounts 2.1.1[41], assigning reads to annotated genes based on the merged GTF.

Downstream analysis was conducted in R. Count matrices were imported into DESeq2 1.46.0[42], where size-factor normalisation, dispersion estimation, and negative binomial modelling were used to quantify differential gene expression between treatments. Standard DESeq2 quality-control steps were applied, including inspection of library size distributions, sample clustering, and principal component analysis. Genes with low counts across (mean < 1 and total count <30) were removed prior to statistical testing. Differential expression results were extracted using DESeq2's Wald test, with multiple-testing correction applied using the Benjamini-Hochberg method.

#### Functional analyses

Functional regions of the myosin gene were determined using ScanProsite (https://prosite.expasy.org/scanprosite/). To establish the specific myosin I isoform, a protein alignment was constructed using reference myosin sequences obtained from elsewhere[43], using only the myosin head domain because it is the only region consistently conserved across all myosin classes. After removing alignment gaps, 419 amino-acid positions were retained for phylogenetic analysis. Sequences were aligned with MAFFT using the L-INS-i algorithm, which is designed for accurate alignment of sequences sharing local homology, with minor manual adjustments to improve alignment quality. A phylogeny was then constructed, using IQ-TREE 3.0.1 with ModelFinder to automatically find the best-fit substitution model, and 1000 bootstrap replicates.[44]

Tests for selection were conducted using the RELAX method[45] implemented on the Datamonkey web server (https://www.datamonkey.org). The aligned myosin gene was uploaded, with the sinistral lineage specified as the *test* branch set. RELAX fits a hierarchical codon model to estimate the selection-intensity parameter K, which quantifies whether selective pressure on the test branches is relaxed ($K < 1$) or intensified ($K > 1$) relative to the reference branches. Statistical significance was assessed using a likelihood-ratio test (LRT) between the null model ($K = 1$) and the alternative model (K estimated). Episodic diversifying selection was tested using BUSTED[46], again on Datamonkey. BUSTED fits a random-effects branch-site codon model that allows ω (dN/dS) to vary across both sites and branches, and compares an unconstrained model permitting $\omega > 1$ on foreground branches with a constrained null model in which $\omega \leq 1$. Statistical significance was assessed using a likelihood-ratio test. The analysis incorporated site-to-site synonymous rate variation to reduce false positives. Only the specified foreground branch was tested for episodic diversifying selection.

To investigate potential functional differences between myosin allelic variants, a dual strategy was used. Protein structure models for the sinistral and dextral myosin variants were generated with SWISS-MODEL[47], a template-based homology modelling method that builds 3D structures using experimentally determined templates from the Protein Data Bank (PDB). SWISS-MODEL performs comparative modelling by identifying structural templates, aligning the query sequence, building a 3D model based on the alignment, and evaluating the model quality. To complement the template-based models, AlphaFold2 predictions for the sinistral and dextral variants were generated using ColabFold[48], an implementation of AlphaFold2 with accelerated MMseqs2-based multiple-sequence alignment generation. Then, predicted models from both outputs were inspected to identify structural contexts surrounding sinistral/dextral variant sites, including proximity to conserved residues, secondary-structure elements, and putative functional domains. Specifically, structural comparisons between sinistral and dextral myosin variants were carried out in UCSF ChimeraX v. 1.11.1[49]. The structures from SWISS MODEL or AlphaFold were superimposed using MatchMaker, which aligns proteins based on sequence and backbone secondary-structure similarity, so producing an RMSD value that quantifies overall structural divergence between variants and enables residue by residue comparison. Then, for each amino acid variable position, local structural effects of individual amino acid substitutions were examined by isolating the residue of interest and analysing hydrogen bonds, steric clashes, and Cα–Cα positional differences, measured with the distance command, as a direct geometric measure of how far the two variants diverge at that position. This approach tested whether each substitution is likely to alter local hydrogen bonding networks, packing geometry, or backbone positioning. Further structural analyses were performed using a structure of actin-bound myosin-1c in the ADP-bound state [50] (PDB ID: 9CFU), with $Mg^{2+}$ in the active site and including calmodulin bound to the lever arm.

Conservation profiles and the biochemical nature of amino-acid changes were assessed to distinguish conservative from non-conservative substitutions. Functional impacts of the variants were further evaluated using PROVEAN[51], which predicts the likelihood that an amino-acid substitution affects protein function based on evolutionary conservation and sequence homology.

Finally, given that some of the changes were shown to be exceptional, as a comparison point, we derived myosin 1a/b sequences from other gastropods, using the MolluscaGenes workflow[52], focussing in particular on two other groups of snails that show variation in chirality, sinistral Planorbids (*Biomphalaria* / *Bulinus*) and a Physid (*Physa*) against dextral *Lymnaea*, all in the Hygrophila[53].

### Supplementary Text

Further detail of selected parts of the results

#### Inheritance of chirality in *Euhadra* and test for pathology

To infer the dominance relations of chirality, maternal genotype was inferred from offspring chirality, using data from four snails (E186, E177, E181, E182) from the Niigata site #4 (see Table 3). This partial genotype information was then compared with the inferred individual haplotypes for the chromosome 27 region. For example, a dextral snail E186 ("dex") produced sinistral offspring, so it must be genetically sinistral ("dex*S*"). Snail E186 has two very divergent haplotypes at the chr. 27 locus, one of which defines dextral and one that defines sinistral. E186 is therefore a dextral snail of genotype *DS*; the *S* allele is dominant to the *D* allele. Consistent with these same inferences, E177 was dex*DD*, E181 was sin*DS* and E182 was sin*SS*.

To confirm that the sinistral or dextral allele does not cause pathology in *E. quaesita*, we compared egg hatch rates. A test for the effect of chirality and sampling location on hatch rate showed neither as significant (chirality: $P = 0.42$, W = 132; sampling location: $P = 0.58$, W = 134.5). Using a generalized linear mixed models (GLMMs), the model excluding the interaction term provided the better fit. Similarly, under the final (additive) model, neither chirality nor sampling location significantly predicted average hatch rate (chirality: estimate = 0.28, SE = 0.34, $P = 0.41$; sampling location: estimate = 0.012, SE = 0.35, $P = 0.96$).

#### *De novo* genome assembly and annotation

A detailed report on the *E. quaesita* genome assembly and annotation will be published elsewhere. In brief, genome assembly was supported by 146 gigabase pairs of HiFi sequence data (~32x depth, 14.5 million reads, N50 = 12.9 kb) and 12.4 Gb of Nanopore long reads (2.8 million reads, N50 = 5.6 kb). The initial Hifiasm assembly produced 10116 contigs totalling 5.3 Gb (contig N50 = 1.12 Mb); after haplotig purging, the assembly was 7,361 contigs totalling 4.5 Gb (contig N50 =1.32 Mb). Hi-C sequencing yielded 1.6 billion paired-end reads (~53x depth). The final chromosome-scale assembly comprised 174 scaffolds totalling 4.54 Gb, with 28 major scaffolds corresponding to putative chromosomes, ranging from 94.6 Mb to 310.5 Mb. The scaffold N50 was 161.0 Mb, with 99.96 percent of the genome contained in scaffolds longer than 50 kb. Contiguity was high, with 7,335 contigs (contig N50 = 1.27 Mb; maximum contig length = 9.11 Mb) and only 0.014 percent gap content. BUSCO analysis (metazoa_odb10) identified 96 percent complete genes (82.5 percent single-copy, 13.5 percent duplicated), with 2.1 percent fragmented and 1.9 percent missing (n = 954), indicating a highly complete assembly.

#### Whole genome re-sequencing, mapping to reference, variant calling and filtering

Following fastp quality control and trimming, reads were generated to ~5-10x depth for most samples (**Table S2**). These sequences were then aligned and variants called. Prior to filtering, we examined the mean depth for each of the unfiltered variants in the vcf file, in other words the number of reads that map to a position, subsequently using a depth filter of between 3 and 10 in vcftools 0.1.16. On an individual level, all but one sample (E15) had a mean depth greater than 3. Three individuals were missing a substantial portion of sites (E15, E195, E239). Site information may be missing from E195 (*Mandarina ponderosa*) and E239 (*E. sadoensis*) because they represent the individuals most deeply diverged from the reference species *E. quaesita*. In subsequent filtering, datasets were produced by varying the required proportion of non-missing genotypes. The most stringent dataset, using a 90% filter, retained ~79000 SNPs. Similarly, the 85% dataset retain 1.85 million SNPs, and the 80% and 70% datasets retained, 7.5 million and 44.7 million SNPs, respectively.

#### Genome wide association

GWAS was performed using a vcf (85% filter) with 1,850,850 SNPs. First, a genomic relationship matrix (GRM) was calculated to account for relatedness and background population structure, and association testing then conducted using a linear mixed model (LMM) with a binary response variable (dextral = 1, sinistral = 0). The GRM random effect effectively controlled for stratification resulting from population subdivision and admixture within the contact zone.

#### Single cell RNA-seq methods

We generated a total of 22 single cell libraries for various *Euhadra* species, and 32 libraries for *L. stagnalis*. For *E. quaesita*, there were samples from six homozygous dextral snails, but with replicate pairs from the same egg batch for some (4 snails x 2 and 2 singletons, n=10 total), three of which were from an allopatric dextral site (O2, O4, O5), and three from the site #4 contact zone (M1, M3, M18). Libraries were also made from five homozygous sinistral snails (3 snails x 2 and two singletons, n=8), with two from allopatric sites (C1, G1) and three from the site #4 contact zone (M8, M22, M54). For qualitative comparisons, we also included one individual (M70) that was a *DS* heterozygote, one pair from another dextral snail species, *E. brandtii* (Eb, n=2), and a singleton from a sinistral snail *E. grata* (Eg). For *L. stagnalis*, libraries were generated from egg batches from 8 dextral and 8 sinistral snails, taking two embryos per egg batch (n=16 dextral, n=16 sinistral).

Per cell sequencing depth ranged from 1.43 to 10.26 million reads (mean=5.14), which was reduced to 0.85 to 7.08 million (mean=3.99) after removing RNA and other repetitive sequences. It was noted that a few individuals retained rather few sequences, specifically a batch that was first processed, including *E. brandtii* (Eb_1, Eb_2), *E. grata* (Eg_1), and *E. quaesita* (M70_1, O2_1). A possible cause is that the vitelline fluid was not removed properly, which negatively impacted library construction. These individuals were not included in the main DEseq analysis. HISAT2 mapped an average of 50% of the *Euhadra* reads (excluding the problematic individuals) and 69% of *L. stagnalis* reads per cell to the respective reference genomes, with only ~0-2% mapping to more than one location.

Raw gene-level count data produced by featureCounts were imported into the DESeq2 package. For *Euhadra*, an initial principal component analysis of all samples, and using the top 500 most variable genes, showed that the five samples (four individuals) with low read count were separate from the others (Figure S3), which could be due to a batch effect, but also because three are a different species one is a heterozygote for chirality.

The same analysis was therefore repeated, but excluding data from these five samples. Pairs of embryos from the same individual tended to cluster together. For *L. stagnalis*, the principal component analysis separated dextrals and sinistral on the first axis, with sinistrals forming a particularly tight cluster.

Embryos from the same mother are not independent replicates, so in all subsequent analyses duplicate experiments (reads from paired embryos derived from a single mother) were collapsed into a combined dataset. The main datasets in all subsequent analyses were therefore five homozygous dextral and five homozygous sinistral *E. quaesita* snails, and eight dextral and eight sinistral *L. stagnalis* snails. Principal component analysis of these datasets showed the same pattern (**Figure S11A**), some separation by chirality for *E.*

*quaesita*, and complete separation for *L. stagnalis*. These data formed the group for differential sequence (DEseq) analysis.

To visualise genome-wide patterns of differential expression, we generated an MA plot in which the mean of the normalised counts for each gene was plotted against the $\log_2$ fold change (sinistral versus dextral; **Figures 2D, S11B**). Genes with an adjusted P-value < 0.05 were highlighted, with those up-regulated in sinistral individuals shown in red and those up-regulated in dextral individuals shown in blue; non-significant genes were plotted in grey. This representation allows the distribution of expression changes to be assessed across the full dynamic range of gene expression. In *E. quaesita*, we detected relatively few differentially expressed genes: 8 were significantly up-regulated in sinistrals and 14 were down-regulated (22/26839=0.08%), with most expression differences occurring among moderately expressed genes (**Tables S6, S7**). Nonetheless, a heatmap of the 50 most strongly differentiated genes separated individuals according to chirality phenotype (**Figure S11C**).

In contrast, in *L. stagnalis,* differential expression was much more extensive, with 1,459 genes up-regulated in sinistrals and 1,546 down-regulated; the corresponding heatmap clearly partitioned individuals into sinistral and dextral clusters (3005/17470= 17%) (**Figures 2, S11; Tables 8,9**). The most strongly differentiated gene was a putative actin-related protein 6 (FDR *P* value 1.66E-65); the gene that causes the chiral variation in *L. stagnalis*, formin *dia2* was the 35$^{th}$ most differentiated (FDR *P* value 2.75E-16), down-expressed, highly expressed in dextrals relative to dia1, and down-expressed in the mutant sinistral, consistent with previous.[54]

### Supplementary figures and tables

#### Figure S1.

**Maximum likelihood (top) and ASTRAL whole genome phylogenies showing relationships with *E. quaesita* and to other *Euhadra* species.** Branches are coloured according to known chirality genotype, either red or blue for sinistral or dextral, respectively. The chirality genotype of some individuals at site #4 in Niigata was not known (prior to the GWAS) because they did not produce offspring, so they are shown in dark grey. The four wholly sinistral *Euhadra* species, *E. scaevola*, *E. grata*, *E. decorata* and *E. murayamai* are part of a monophyletic group that also contains the chirally variable *E. quaesita*. Within *E. quaesita*, samples cluster according to the four main groups from sites #1 to #4. The progressively longer branch is probably an artefact, arising from alignment of all species to *E. quaesita* genome assembly. Branch lengths are given as nucleotide substitution rate (ML) and coalescent units (ASTRAL).

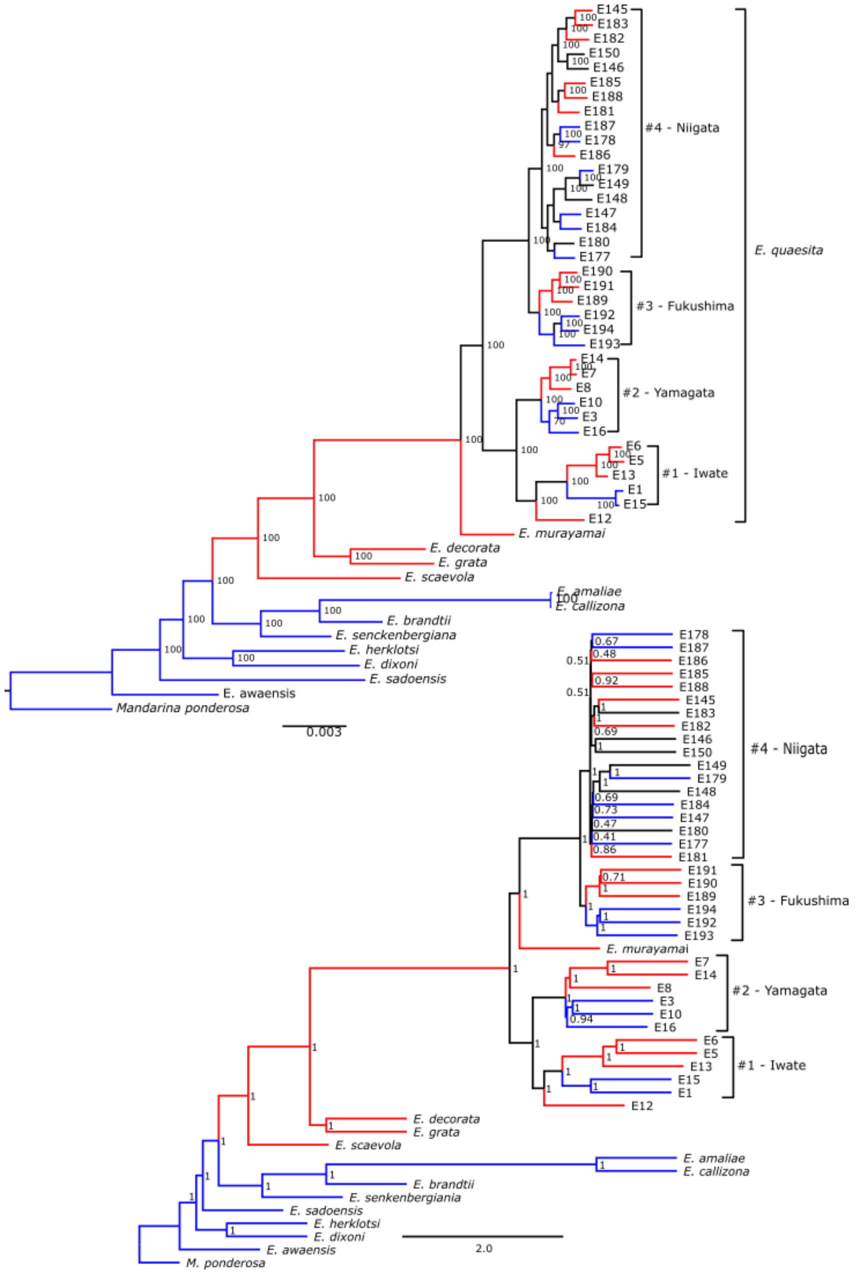

#### Figure S2.

**Admixture plots for *E. quaesita* samples from the four contact zones.** K=2 is optimal number of populations; K=4 separates snails according to geography.

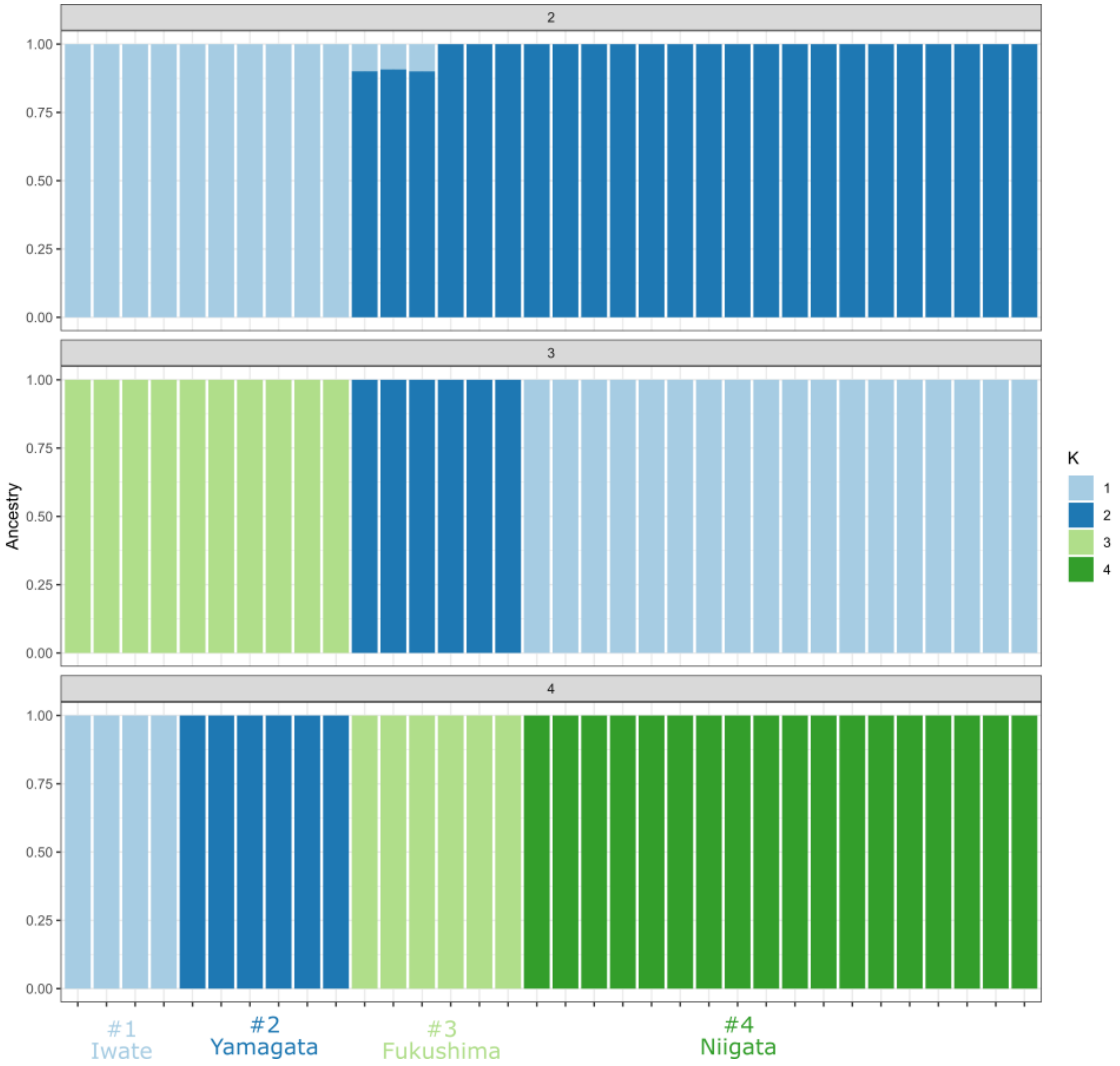

### Figure S3.

Principal components analysis of genomic variation in *E. quaesita*. Most axes of variation tend to separate individuals by geography rather than chirality.

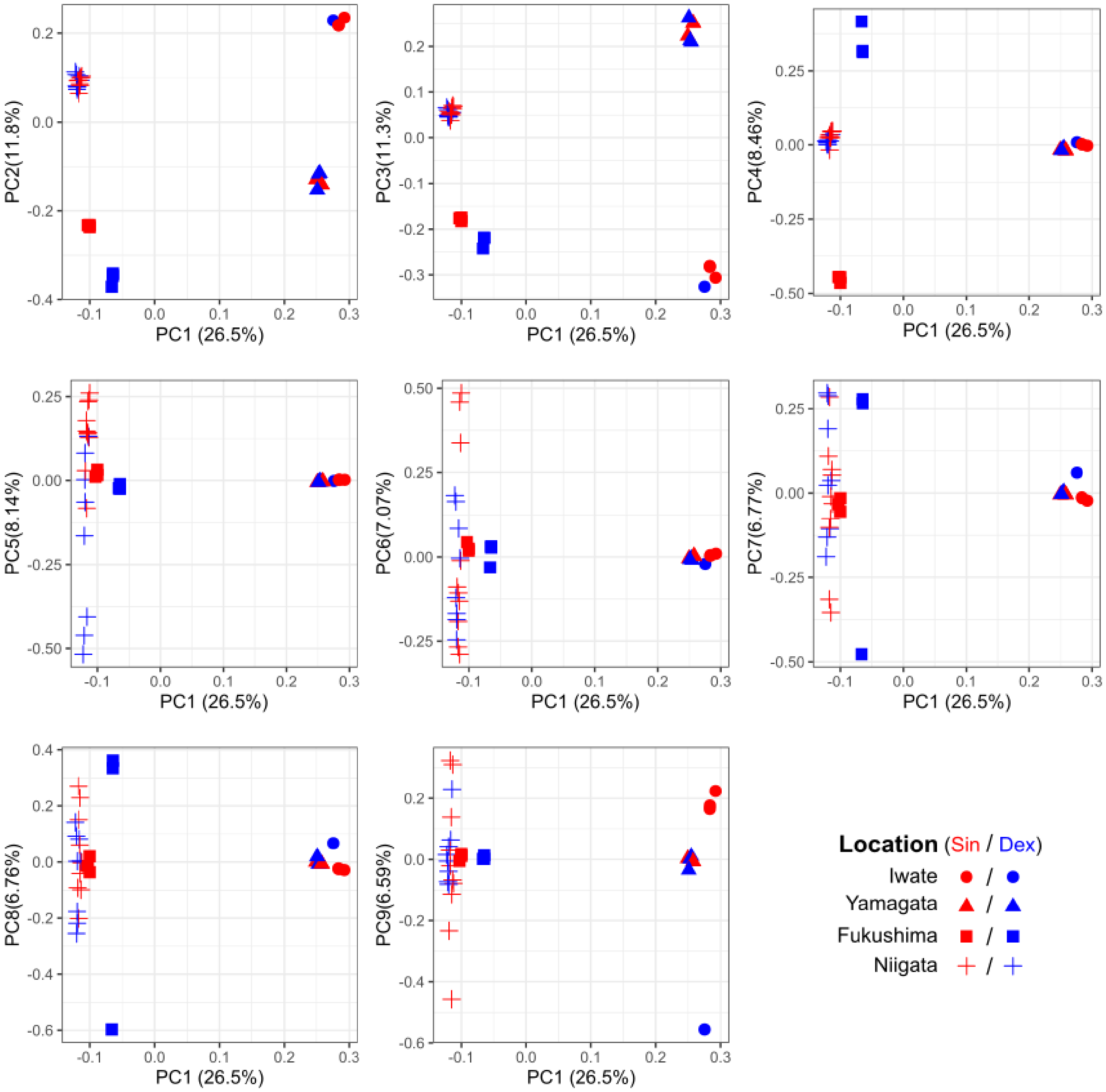

### Figure S4.

**Pairwise Fst and Dxy comparisons between dextral and sinistral snails at three contact zones.** A region of chromosome 27 shows high Fst in all three comparisons (red arrow), whereas the region on chromosome 4 is not differentiated at the Yamagata site. Dxy is does not show any major regions of differentiation in any comparison. Comparisons were not made at site #1 because there were insufficient dextral individuals to be able to phase haplotypes. Red lines show 99th and 99.9th percentiles, black lines show genome-wide mean.

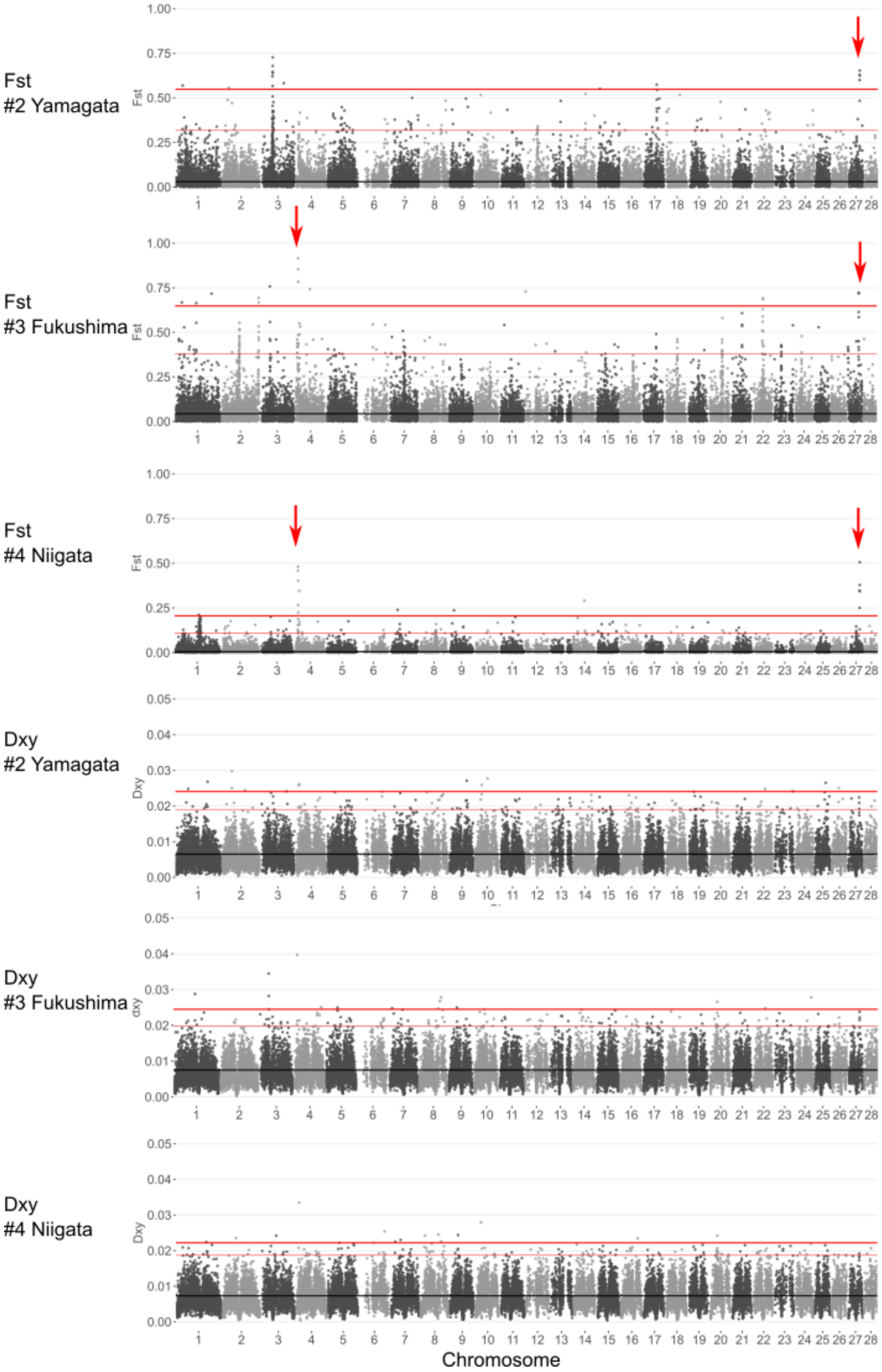

#### Figure S5.

Quantile-quantile (QQ) plot of GWAS results. The diagonal line represents the expected distribution of test statistics, with the deviation from this line at the right hand likely due to an excess of low *P* values, consistent with a true genetic associations.

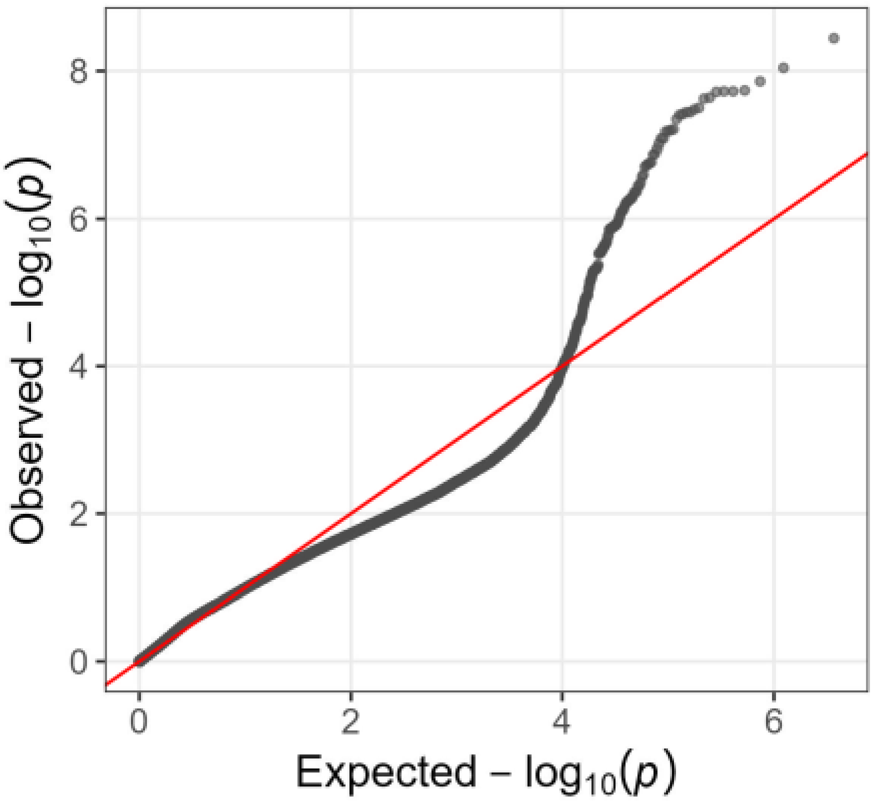

### Figure S6.

LD decay across each of the chromosomes in *E. quaesita*.

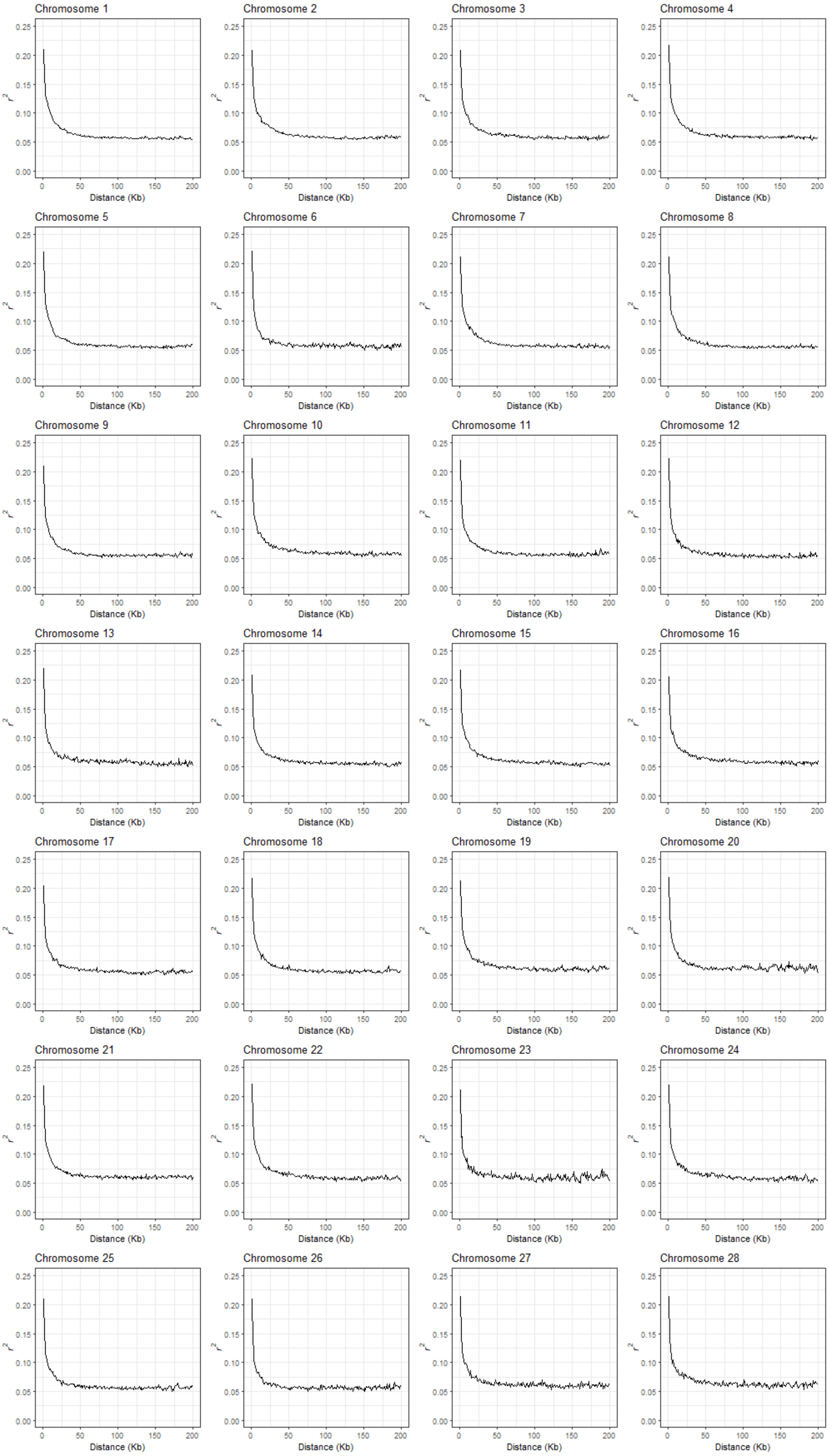

### Figure S7.

**Topology weighting analysis between pairs of populations.** A topology weighting analysis was carried out for two dextral versus two sinistral populations, Yamagata and Fukushima, across each of the chromosomes. Three possible topologies are possible for which only one has the same chiral types clustering together. Shown are the only two regions of the genome that have a region that groups populations by chirality. On chromosome 4, six contiguous non-overlapping windows (defined by 50 sites) cluster by chirality, and on chromosome 27, 139 windows contiguous cluster by chirality.

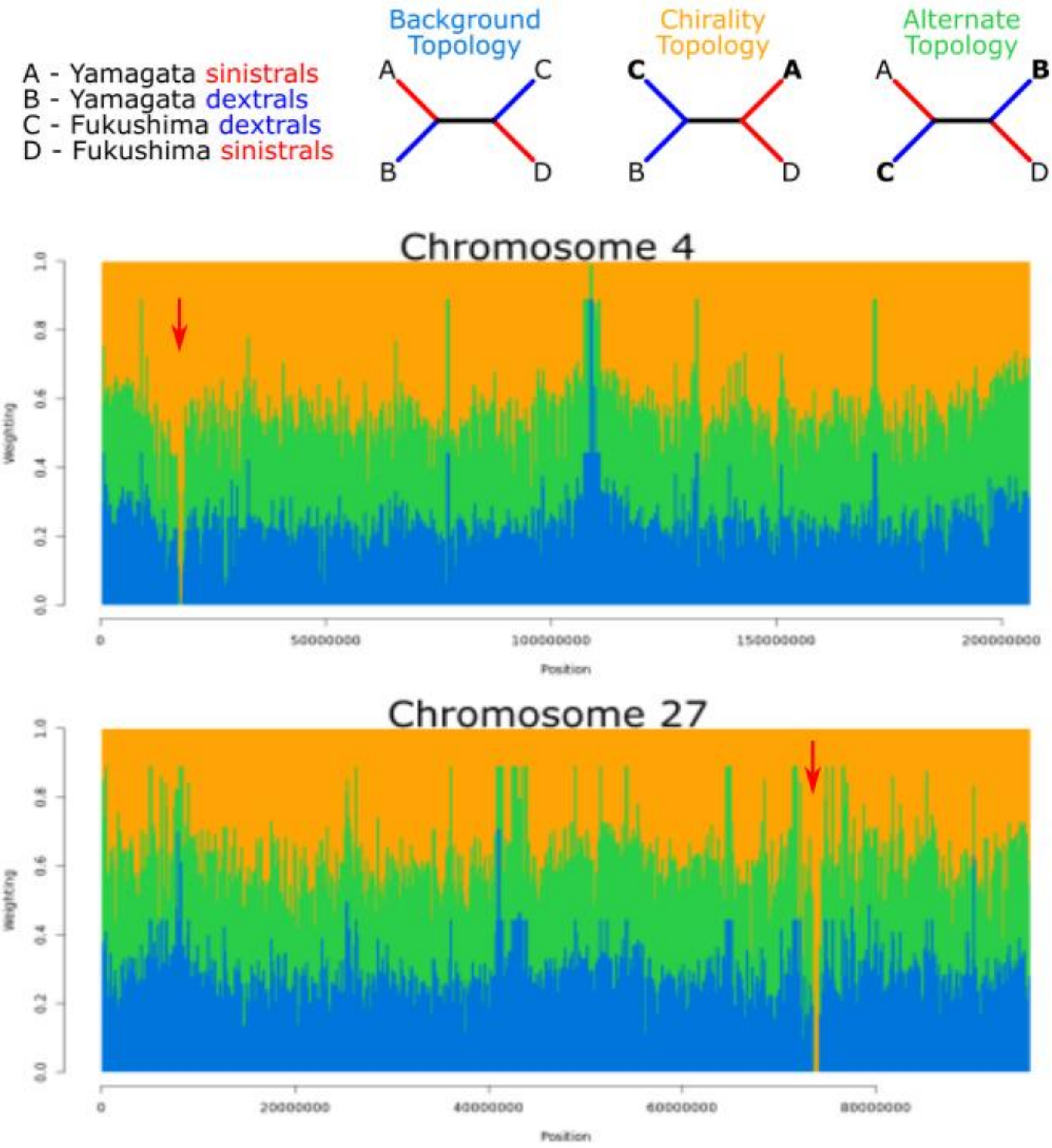

### Figure S8.

**Phylogenies showing relationships between *E. quaesita*, using individual haplotypes.** The phylogeny for the chromosome 27 region perfectly separates dextral (blue) from sinistral (red), the only region to do so in the whole genome; individuals that are heterozygotes for the two lineages are highlighted, all are from the #4 Niigata site. In comparison, for the chromosome 4 region two individuals are anomalous (both homozygotes, shown by red arrows). Individual haplotypes were only produced for *E. quaesita*, so no other species are shown

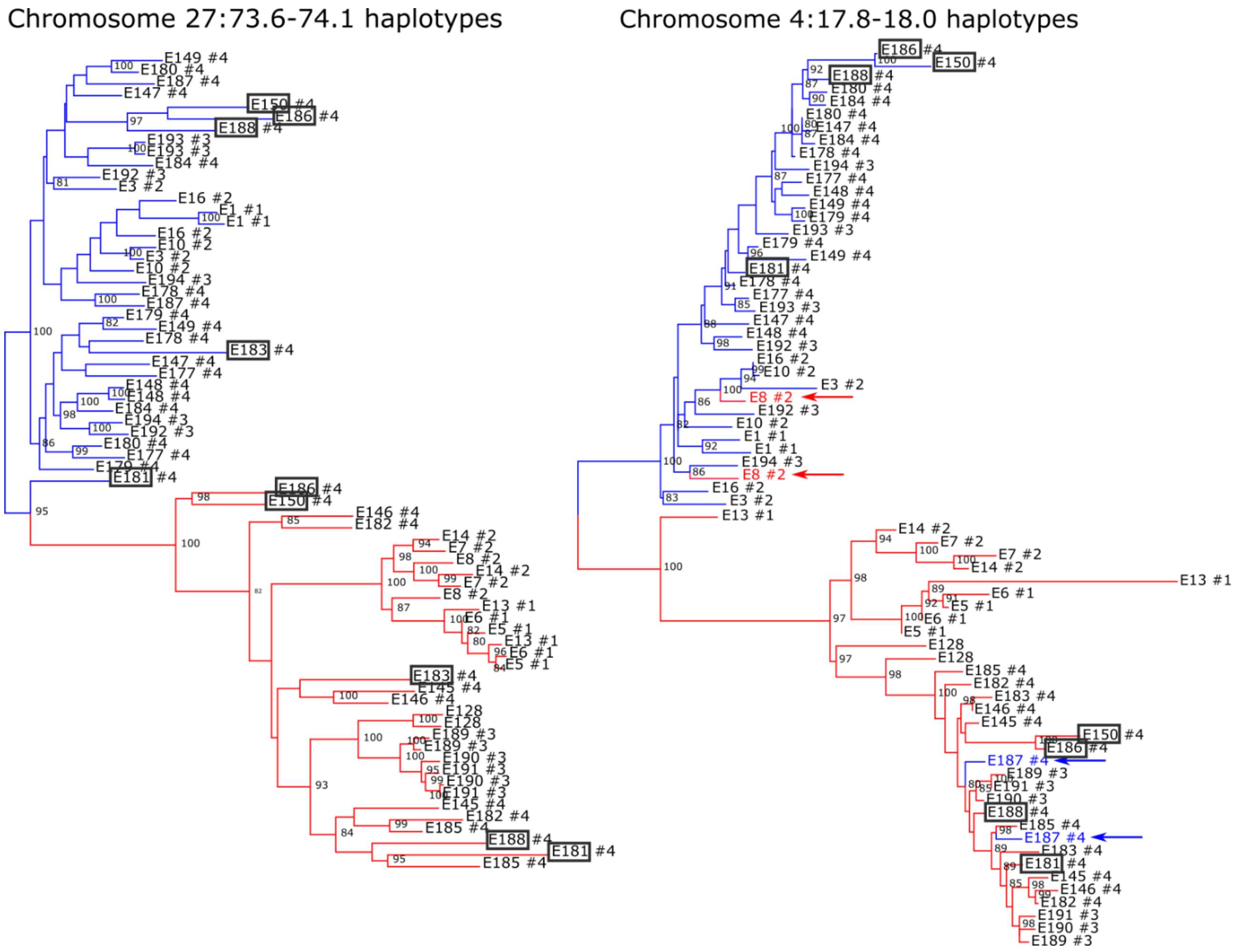

### Figure S9.

**Independent dataset was used to map the chirality locus.** GWAS was carried out on ddRAD-seq derived data, using 107 samples of *E. quaesita*. Mean per-sample coverage was 12.7 (S.D. = 5.7), retaining 84,756 loci. The results are in line with the main GWAS analysis in that there are two major peaks that are associated with chirality, on chromosomes 27 and 4. The Q-Q plot shows some limited inflation of the P values.

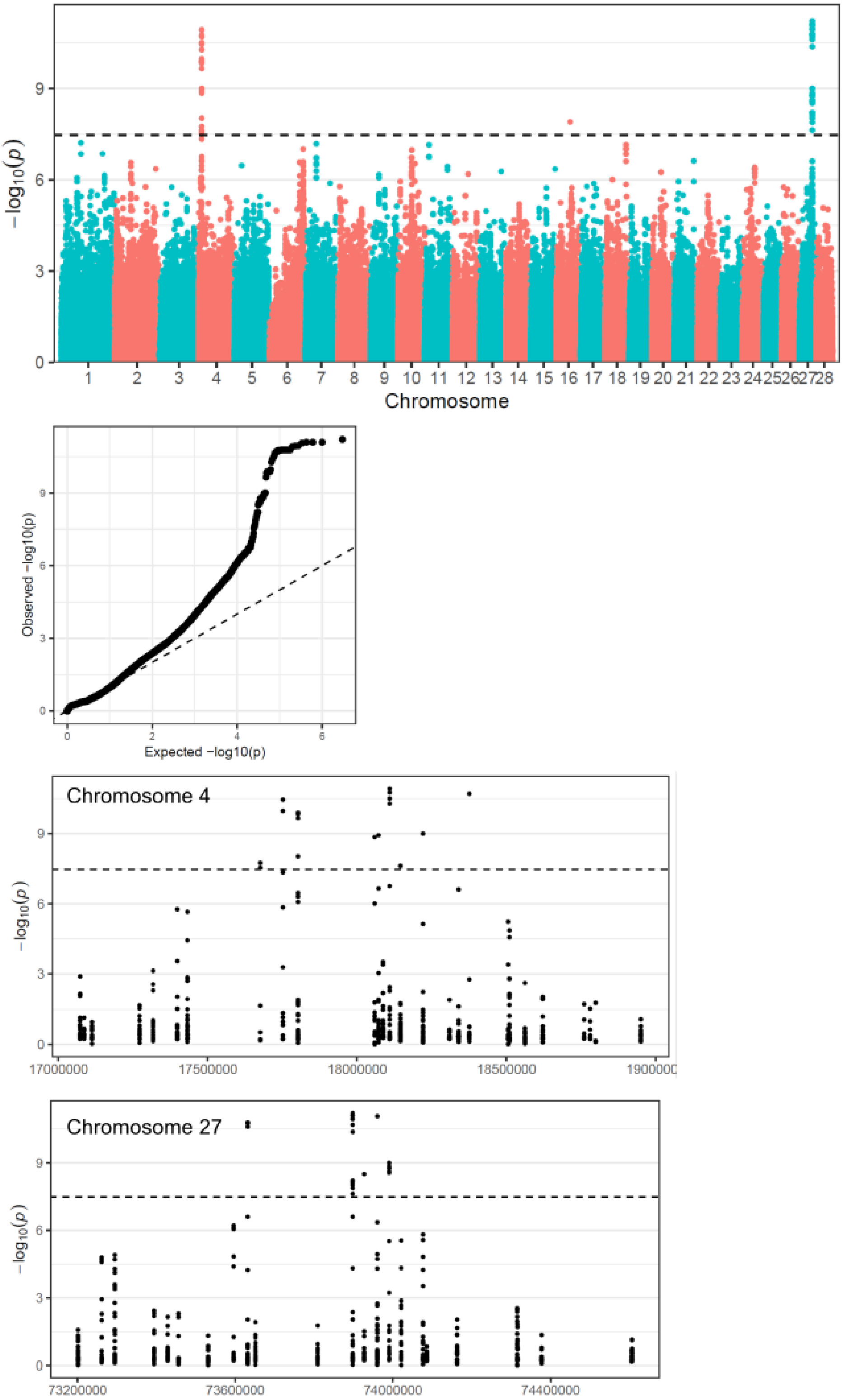

### Figure S10.

**Phylogenies showing relationships between *E. quaesita* and all other species.** The phylogeny for the chromosome 27 region perfectly separates all dextral species and individuals (blue) from all sinistral species and individuals (red). In comparison, two individuals are anomalous for the chromosome 4 region, E8 and E187.

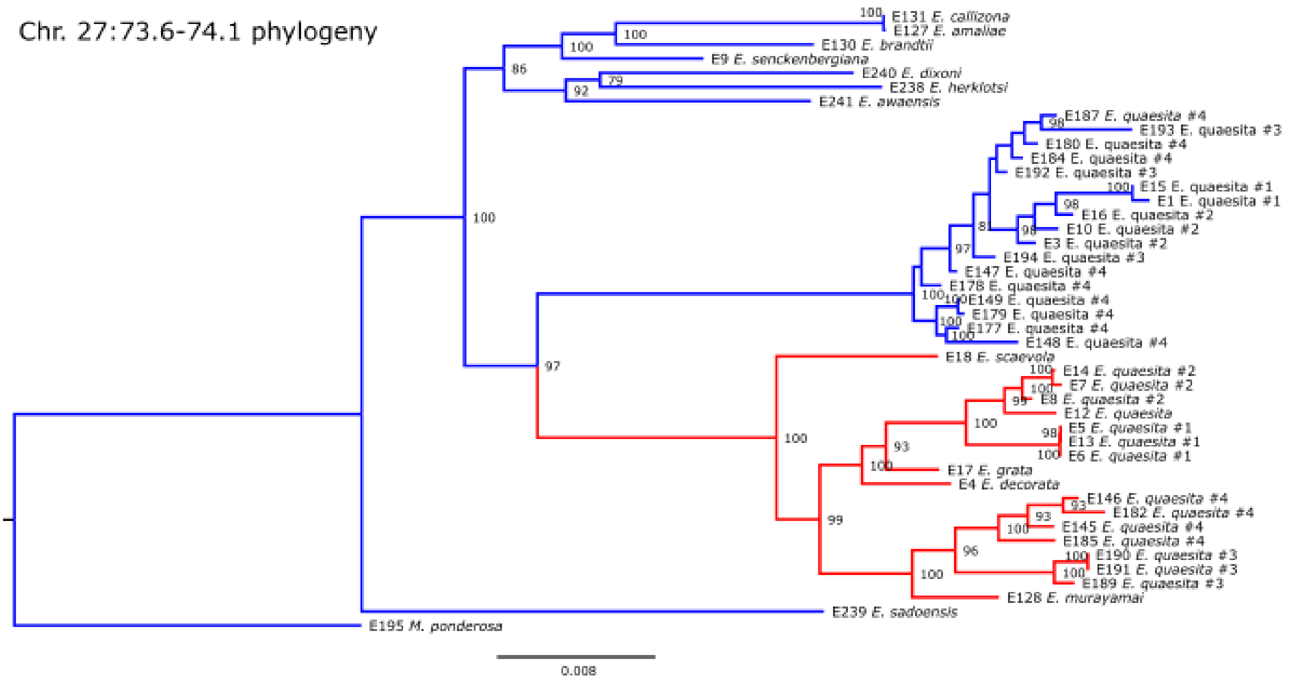


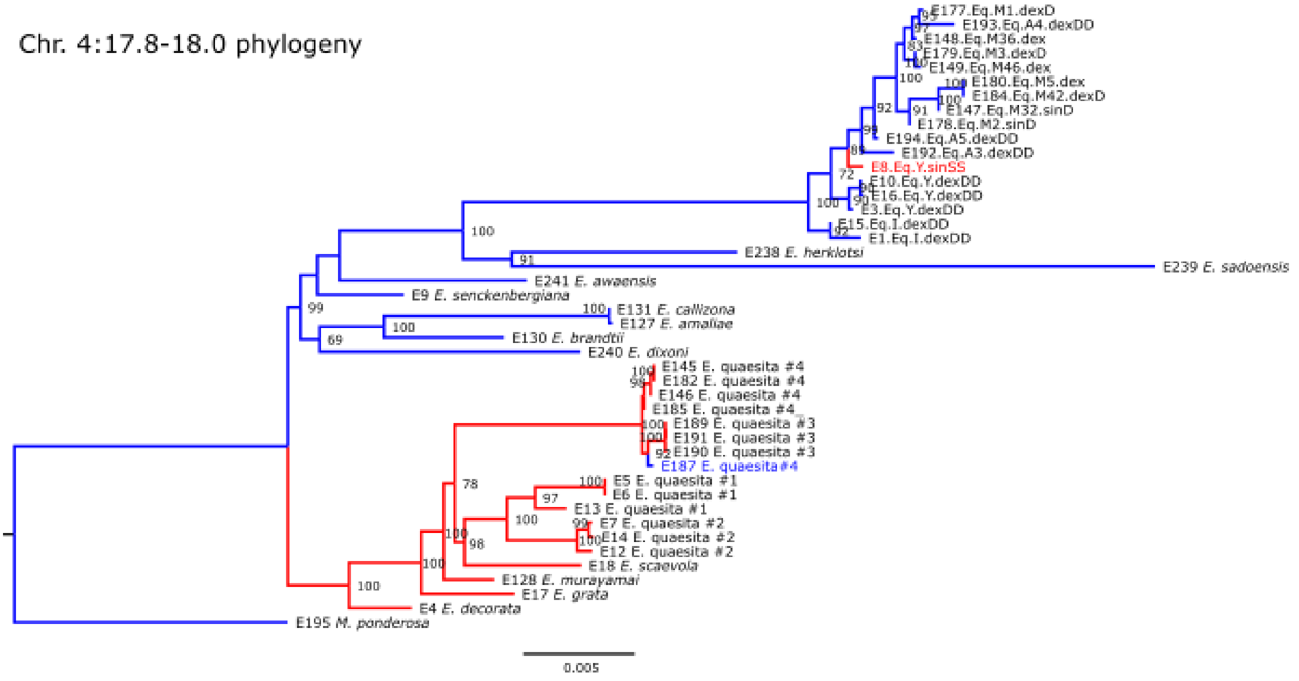

### Figure S11.

**RNAseq analysis of single-cell embryos of *E. quaesita* (left) and *L. stagnalis.* a,** Principal component analysis (PCA) of normalised gene expression values showing relationships among individuals based on genome-wide expression patterns. Samples are coloured by chirality phenotype (red: sinistral versus blue: dextral); axes indicate the percentage of variance explained by each principal component. Sinistral and dextral *E. quaesita* do not separate on the first two axes, unlike *L. stagnalis*. **b,** Volcano plot showing log-fold change (sinistral versus dextral) plotted against log adjusted *P* value for all genes. Genes with an adjusted *P* value < 0.05 are highlighted, with significant up-regulation in sinistrals shown in red and significant down-regulation shown in blue. *E. quaesita* have very few genes that are differentially expressed compared with *L. stagnalis*. **c,** Heat map of the 50 genes showing the strongest differential expression between sinistral and dextral individuals. Hierarchical clustering of genes and samples reveals separation of individuals according to chirality phenotype.

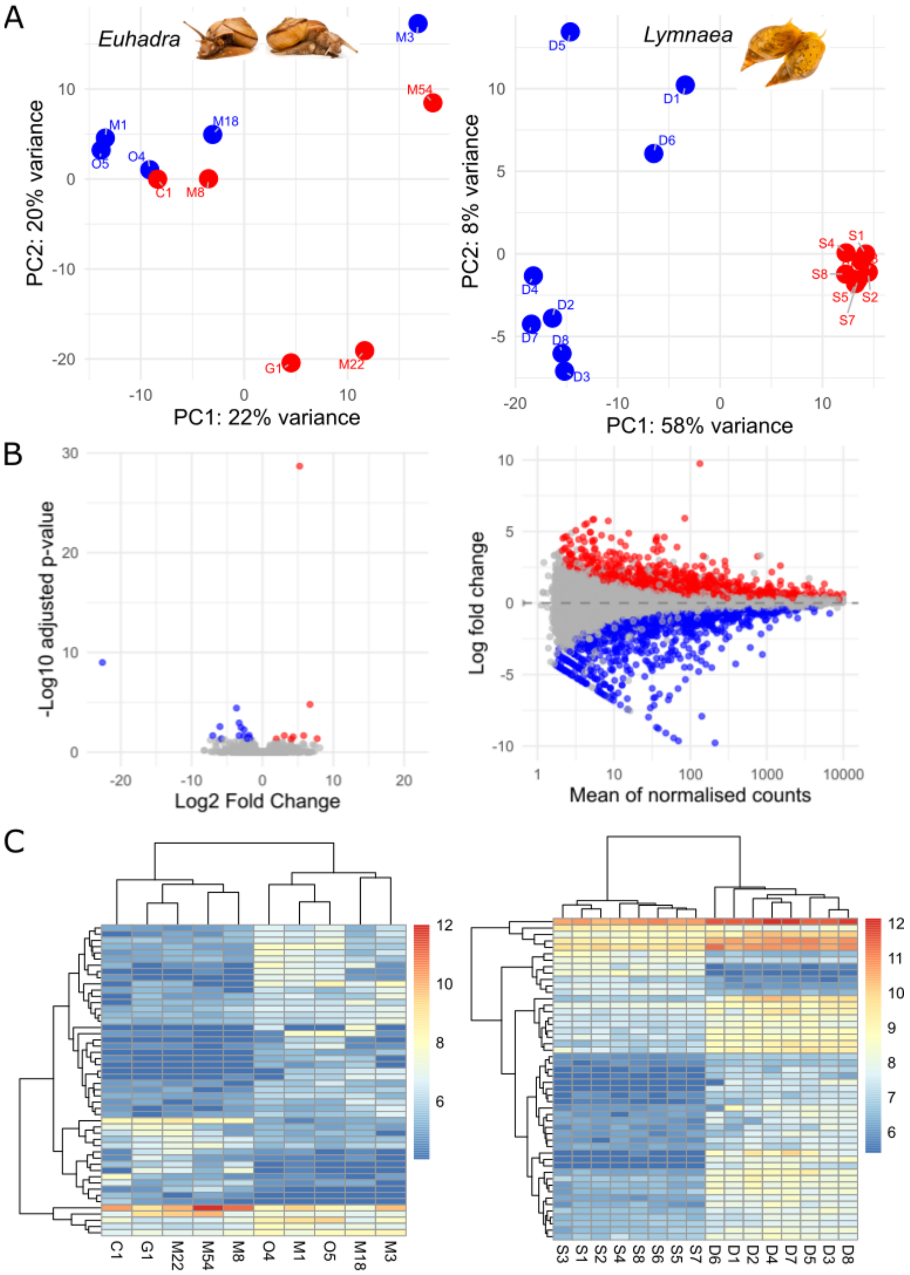

### Supplementary Videos

#### Video S1.

Time-lapse showing dextral twist of *E. quaesita* embryo during third cleavage.

#### Video S2.

Video showing courtship and unilateral mating between sinistral and dextral *E. quaesita*. Speed is 4x normal. https://youtu.be/GvXJtb9aesA

### Table legends

**Table S1. Summary of samples used for genomic analyses.**

**Table S2. Samples used, including source population, chirality information and basic genome sequencing statistics.**

Chirality genotype is assumed to correspond to shell phenotype (e.g. sinistral snail is genotype *SS*) when species or populations are invariant for chirality. For populations that vary in chirality (#4 Niigata only), maternal chirality genotype (a single *S* or *D* allele) was inferred by recording the chirality of offspring from the mother; "maternal genotype" is the character that was used for the GWAS. Following the genomic analyses, we were able to infer that sinistral is dominant and thus the full chirality genotype of individual snails, noted as "inferred chirality genotype (GWAS)". This was achieved by inspecting the individual haplotypes present in the chromosome 27 region alongside the offspring chirality (Figure 3A; Table 3).

**Table S3. Summary showing hatch rate of egg batches from sinistral and dextral *E. quaesita*.**

Location #4 in Niigata is where there is present-day gene-flow between the two chiral types. Maternal phenotype is the chirality of the mother's shell. Maternal genotype is the chirality genotype of the mother, inferred by the shell chirality of her offspring. Inferred chirality genotype (GWAS) is the full chirality genotype, inferred by the chirality of offspring and/or genotype in the chromosome 27 candidate region.

**Table S4. SNPs associated with chirality by GWAS.**

SNPs shown in bold are also significantly associated with chirality using the more conservative Bonferroni correction.

**Table S5. Genes on chromosome 27 and chromosome 4 candidate regions.**

Details includes whether gene expressed in single-cell embryo, and also whether local gene region phylogeny is congruent with chirality. Myosin is the only gene that showed a significant association in the GWAS, is expressed in the single cell embryo and has a congruent phylogeny. None of the genes showed significant differences in expression in the single-cell embryo.

**Table S6. Differentially expressed genes in *E. quaesita.***

Genes associated with differential expression in *E. quaesita* single-cell embryos

**Table S7. Normalised expression data for all transcripts from *E. quaesita* single-cell embryos.**

**Table S8. Differentially expressed genes in *L. stagnalis*.** Genes associated with differential expression in *L. stagnalis* single-cell embryos

**Table S9. Normalised expression data for all transcripts from *L. stagnalis* single-cell embryos.**

**Table S10. Functional variation between dextral and sinistral versions of the myosin gene.**

**Table S11. Individuals of *E. quaesita* used for ddRAD-seq GWAS.**

### References


1. Ueshima, R., and Asami, T. (2003). Single-gene speciation by left-right reversal - A land-snail species of polyphyletic origin results from chirality constraints on mating. Nature *425*, 679-679.
2. Richards, P.M., Morii, Y., Kimura, K., Hirano, T., Chiba, S., and Davison, A. (2017). Single-gene speciation: Mating and gene flow between mirror-image snails. Evolution Letters *1*, 282–291. 10.1002/evl3.31.
3. Davison, A., McDowell, G.S., Holden, J.M., Johnson, H.F., Koutsovoulos, G.D., Liu, M.M., Hulpiau, P., Van Roy, F., Wade, C.M., Banerjee, R., et al. (2016). Formin is associated with left-right asymmetry in the pond snail and the frog. Current Biology *26*, 654-660. 10.1016/j.cub.2015.12.071.
4. Davison, A. (2020). Flipping shells: unwinding LR asymmetry in mirror-image molluscs. Trends in Genetics *36*, 189-202. 10.1016/j.tig.2019.12.003.
5. Davison, A., Barton, N.H., and Clarke, B. (2009). The effect of coil phenotypes and genotypes on the fecundity and viability of *Partula suturalis* and *Lymnaea stagnalis*: implications for the evolution of sinistral snails. Journal of Evolutionary Biology *22*, 1624-1635. 10.1111/j.1420-9101.2009.01770.x.
6. Noda, T., Satoh, N., and Asami, T. (2019). Heterochirality results from reduction of maternal diaph expression in a terrestrial pulmonate snail. Zoological Letters *5*, 2, 2. 10.1186/s40851-018-0120-0.
7. Cheng, H., Jarvis, E.D., Fedrigo, O., Koepfli, K.-P., Urban, L., Gemmell, N.J., and Li, H. (2022). Haplotype-resolved assembly of diploid genomes without parental data. Nat. Biotechnol. *40*, 1332-1335. 10.1038/s41587-022-01261-x.
8. Guan, D., McCarthy, S.A., Wood, J., Howe, K., Wang, Y., and Durbin, R. (2020). Identifying and removing haplotypic duplication in primary genome assemblies. Bioinformatics *36*, 2896-2898. 10.1093/bioinformatics/btaa025.
9. Li, H., and Durbin, R. (2009). Fast and accurate short read alignment with Burrows–Wheeler transform. Bioinformatics *25*, 1754-1760. 10.1093/bioinformatics/btp324.
10. Danecek, P., Bonfield, J.K., Liddle, J., Marshall, J., Ohan, V., Pollard, M.O., Whitwham, A., Keane, T., McCarthy, S.A., Davies, R.M., and Li, H. (2021). Twelve years of SAMtools and BCFtools. GigaScience *10*, giab008. 10.1093/gigascience/giab008.
11. Zhou, C., McCarthy, S.A., and Durbin, R. (2023). YaHS: yet another Hi-C scaffolding tool. Bioinformatics *39*, btac808. 10.1093/bioinformatics/btac808.
12. Zeng, X., Yi, Z., Zhang, X., Du, Y., Li, Y., Zhou, Z., Chen, S., Zhao, H., Yang, S., Wang, Y., and Chen, G. (2024). Chromosome-level scaffolding of haplotype-resolved assemblies using Hi-C data without reference genomes. Nat Plants *10*, 1184-1200. 10.1038/s41477-024-01755-3.
13. Durand, N.C., Shamim, M.S., Machol, I., Rao, S.S., Huntley, M.H., Lander, E.S., and Aiden, E.L. (2016). Juicer Provides a One-Click System for Analyzing Loop-Resolution Hi-C Experiments. Cell Syst *3*, 95-98. 10.1016/j.cels.2016.07.002.
14. Robinson, J.T., Turner, D., Durand, N.C., Thorvaldsdóttir, H., Mesirov, J.P., and Aiden, E.L. (2018). Juicebox.js Provides a Cloud-Based Visualization System for Hi-C Data. Cell Syst *6*, 256-258.e251. 10.1016/j.cels.2018.01.001.
15. Flynn, J.M., Hubley, R., Goubert, C., Rosen, J., Clark, A.G., Feschotte, C., and Smit, A.F. (2020). RepeatModeler2 for automated genomic discovery of

transposable element families. Proceedings of the National Academy of Sciences *117*, 9451-9457. doi:10.1073/pnas.1921046117.
16. Lerat, E., Fablet, M., Modolo, L., Lopez-Maestre, H., and Vieira, C. (2017). TEtools facilitates big data expression analysis of transposable elements and reveals an antagonism between their activity and that of piRNA genes. Nucleic Acids Res *45*, e17. 10.1093/nar/gkw953.
17. Gabriel, L., Brůna, T., Hoff, K.J., Ebel, M., Lomsadze, A., Borodovsky, M., and Stanke, M. (2024). BRAKER3: Fully automated genome annotation using RNA-seq and protein evidence with GeneMark-ETP, AUGUSTUS, and TSEBRA. Genome Res. *34*, 769-777. 10.1101/gr.278090.123.
18. Kuznetsov, D., Tegenfeldt, F., Manni, M., Seppey, M., Berkeley, M., Kriventseva, E.V., and Zdobnov, E.M. (2023). OrthoDB v11: annotation of orthologs in the widest sampling of organismal diversity. Nucleic Acids Res *51*, D445-d451. 10.1093/nar/gkac998.
19. Manni, M., Berkeley, M.R., Seppey, M., and Zdobnov, E.M. (2021). BUSCO: Assessing Genomic Data Quality and Beyond. Current Protocols *1*, e323. https://doi.org/10.1002/cpz1.323.
20. Chen, S., Zhou, Y., Chen, Y., and Gu, J. (2018). fastp: an ultra-fast all-in-one FASTQ preprocessor. Bioinformatics *34*, i884-i890. 10.1093/bioinformatics/bty560.
21. Broad Institute (2019). Picard Toolkit. Broad Institute, GitHub repository.
22. Danecek, P., Auton, A., Abecasis, G., Albers, C.A., Banks, E., DePristo, M.A., Handsaker, R.E., Lunter, G., Marth, G.T., Sherry, S.T., et al. (2011). The variant call format and VCFtools. Bioinformatics *27*, 2156-2158. 10.1093/bioinformatics/btr330.
23. Davison, A., Chiba, S., Barton, N.H., and Clarke, B.C. (2005). Speciation and gene flow between snails of opposite chirality. Public Library of Science Biology *3*, e282.
24. Perdry, H., and Dandine-Roulland, C. (2023). gaston: Genetic Data Handling (QC, GRM, LD, PCA) & Linear Mixed Models. R package version 1.6. .
25. Martin, S.H., and Van Belleghem, S.M. (2017). Exploring Evolutionary Relationships Across the Genome Using Topology Weighting. Genetics *206*, 429-438. 10.1534/genetics.116.194720.
26. Ortiz, E.M. (2019). vcf2phylip v2.0: convert a VCF matrix into several matrix formats for phylogenetic analysis. (v2.0). .
27. Minh, B.Q., Schmidt, H.A., Chernomor, O., Schrempf, D., Woodhams, M.D., von Haeseler, A., and Lanfear, R. (2020). IQ-TREE 2: New Models and Efficient Methods for Phylogenetic Inference in the Genomic Era. Molecular Biology and Evolution *37*, 1530-1534. 10.1093/molbev/msaa015.
28. Rabiee, M., Sayyari, E., and Mirarab, S. (2019). Multi-allele species reconstruction using ASTRAL. Molecular Phylogenetics and Evolution *130*, 286-296. https://doi.org/10.1016/j.ympev.2018.10.033.
29. Van Belleghem, S.M., Rastas, P., Papanicolaou, A., Martin, S.H., Arias, C.F., Supple, M.A., Hanly, J.J., Mallet, J., Lewis, J.J., Hines, H.M., et al. (2017). Complex modular architecture around a simple toolkit of wing pattern genes. Nature Ecology & Evolution *1*, 0052. 10.1038/s41559-016-0052.
30. Chang, C.C., Chow, C.C., Tellier, L.C., Vattikuti, S., Purcell, S.M., and Lee, J.J. (2015). Second-generation PLINK: rising to the challenge of larger and richer datasets. GigaScience *4*. 10.1186/s13742-015-0047-8.

31. Alexander, D.H., and Lange, K. (2011). Enhancements to the ADMIXTURE algorithm for individual ancestry estimation. BMC Bioinformatics *12*, 246. 10.1186/1471-2105-12-246.
32. Peterson, B.K., Weber, J.N., Kay, E.H., Fisher, H.S., and Hoekstra, H.E. (2012). Double Digest RADseq: An Inexpensive Method for De Novo SNP Discovery and Genotyping in Model and Non-Model Species. PLoS One *7*, e37135. 10.1371/journal.pone.0037135.
33. Ishii, Y., Ito, S., Kameda, Y., Takano, T., Waki, T., Chiba, S., and Hirano, T. (2026). Reticulate evolution in terrestrial snails of Euhadra peliomphala species complex. Molecular Phylogenetics and Evolution *214*, 108437. https://doi.org/10.1016/j.ympev.2025.108437.
34. Yamazaki, D., Ito, S., Miura, O., Sasaki, T., and Chiba, S. (2022). High-throughput SNPs dataset reveal restricted population connectivity of marine gastropod within the narrow distribution range of peripheral oceanic islands. Sci. Rep. *12*, 2119. 10.1038/s41598-022-05026-z.
35. Eaton, D.A.R., and Overcast, I. (2020). ipyrad: Interactive assembly and analysis of RADseq datasets. Bioinformatics *36*, 2592-2594. 10.1093/bioinformatics/btz966.
36. Catchen, J., Hohenlohe, P.A., Bassham, S., Amores, A., and Cresko, W.A. (2013). Stacks: an analysis tool set for population genomics. Molecular Ecology *22*, 3124-3140. 10.1111/mec.12354.
37. Hagemann-Jensen, M., Ziegenhain, C., Chen, P., Ramsköld, D., Hendriks, G.-J., Larsson, A.J.M., Faridani, O.R., and Sandberg, R. (2020). Single-cell RNA counting at allele and isoform resolution using Smart-seq3. Nat. Biotechnol. *38*, 708-714. 10.1038/s41587-020-0497-0.
38. Kopylova, E., Noé, L., and Touzet, H. (2012). SortMeRNA: fast and accurate filtering of ribosomal RNAs in metatranscriptomic data. Bioinformatics *28*, 3211-3217. 10.1093/bioinformatics/bts611.
39. Kim, D., Paggi, J.M., Park, C., Bennett, C., and Salzberg, S.L. (2019). Graph-based genome alignment and genotyping with HISAT2 and HISAT-genotype. Nat. Biotechnol. *37*, 907-915. 10.1038/s41587-019-0201-4.
40. Pertea, M., Pertea, G.M., Antonescu, C.M., Chang, T.C., Mendell, J.T., and Salzberg, S.L. (2015). StringTie enables improved reconstruction of a transcriptome from RNA-seq reads. Nat. Biotechnol. *33*, 290-295. 10.1038/nbt.3122.
41. Liao, Y., Smyth, G.K., and Shi, W. (2013). featureCounts: an efficient general purpose program for assigning sequence reads to genomic features. Bioinformatics *30*, 923-930. 10.1093/bioinformatics/btt656.
42. Love, M.I., Huber, W., and Anders, S. (2014). Moderated estimation of fold change and dispersion for RNA-seq data with DESeq2. Genome Biol *15*, 550. 10.1186/s13059-014-0550-8.
43. Sebé-Pedrós, A., Grau-Bové, X., Richards, T.A., and Ruiz-Trillo, I. (2014). Evolution and Classification of Myosins, a Paneukaryotic Whole-Genome Approach. Genome Biol. Evol. *6*, 290-305. 10.1093/gbe/evu013.
44. Wong, T.K.F., Ly-Trong, N., Ren, H., Banos, H., Roger, A.J., Susko, E., Bielow, C., De Maio, N., Goldman, N., Hahn, M.W., et al. (2025). IQ-TREE 3: Phylogenomic Inference Software using Complex Evolutionary Models.
45. Wertheim, J.O., Murrell, B., Smith, M.D., Kosakovsky Pond, S.L., and Scheffler, K. (2015). RELAX: detecting relaxed selection in a phylogenetic framework. Mol Biol Evol *32*, 820-832. 10.1093/molbev/msu400.

46. Murrell, B., Weaver, S., Smith, M.D., Wertheim, J.O., Murrell, S., Aylward, A., Eren, K., Pollner, T., Martin, D.P., Smith, D.M., et al. (2015). Gene-wide identification of episodic selection. Mol Biol Evol *32*, 1365-1371. 10.1093/molbev/msv035.
47. Waterhouse, A., Bertoni, M., Bienert, S., Studer, G., Tauriello, G., Gumienny, R., Heer, F.T., de Beer, T.A.P., Rempfer, C., Bordoli, L., et al. (2018). SWISS-MODEL: homology modelling of protein structures and complexes. Nucleic Acids Res *46*, W296-w303. 10.1093/nar/gky427.
48. Mirdita, M., Schütze, K., Moriwaki, Y., Heo, L., Ovchinnikov, S., and Steinegger, M. (2022). ColabFold: making protein folding accessible to all. Nat. Methods *19*, 679-682. 10.1038/s41592-022-01488-1.
49. Meng, E.C., Goddard, T.D., Pettersen, E.F., Couch, G.S., Pearson, Z.J., Morris, J.H., and Ferrin, T.E. (2023). UCSF ChimeraX: Tools for structure building and analysis. Protein Sci. *32*, e4792. https://doi.org/10.1002/pro.4792.
50. Chavali, S.S., Carman, P.J., Shuman, H., Ostap, E.M., and Sindelar, C.V. (2025). High-resolution structures of Myosin-IC reveal a unique actin-binding orientation, ADP release pathway, and power stroke trajectory. Proceedings of the National Academy of Sciences *122*, e2415457122. doi:10.1073/pnas.2415457122.
51. Choi, Y., and Chan, A.P. (2015). PROVEAN web server: a tool to predict the functional effect of amino acid substitutions and indels. Bioinformatics *31*, 2745-2747. 10.1093/bioinformatics/btv195.
52. Pérez-Moreno, J.L., and Katz, P.S. (2026). MolluscaGenes: A transcriptomic database for the Mollusca. bioRxiv, 2026.2005.2005.723003. 10.64898/2026.05.05.723003.
53. Saadi, A.J., Davison, A., and Wade, C.M. (2020). Molecular phylogeny of the freshwater snails and limpets (Panpulmonata: Hygrophila). Zoological Journal of the Linnean Society *190*, 518–531.
54. Johnson, H.F., and Davison, A. (2019). A new set of endogenous control genes for use in quantitative real-time PCR experiments show that formin Ldia2dex transcripts are enriched in the early embryo of the pond snail *Lymnaea stagnalis* (Panpulmonata). Journal of Molluscan Studies *85*, 389–397. 10.1101/660381.
